# Crosstalk between the plasma membrane and cell-cell adhesion maintains epithelial identity for correct polarised cell divisions

**DOI:** 10.1101/2023.05.19.541433

**Authors:** Manal M. Hosawi, Jiaoqi Cheng, Maria Fankhaenel, Marcin R. Przewloka, Salah Elias

**Affiliations:** School of Biological Sciences, University of Southampton, Southampton, SO17 1BJ, United Kingdom; Institute for Life Sciences, University of Southampton, Southampton, SO17 1BJ, United Kingdom

## Abstract

Polarised epithelial cell divisions represent a fundamental mechanism for tissue maintenance and morphogenesis; their dysregulation often leads to developmental disorders and cancer. Morphological and mechanical changes in the plasma membrane influence the organisation and crosstalk of microtubules and actin at the cell cortex, thereby regulating the mitotic spindle machinery and chromosome segregation. Yet, the precise mechanisms linking plasma membrane remodelling to cell polarity and cortical cytoskeleton dynamics to ensure accurate execution of mitosis in mammalian epithelial cells remain poorly understood. Here we experimentally manipulated the density of mammary epithelial cells in culture, which led to several mitotic defects. We found that perturbation of cell-cell adhesion integrity impairs the dynamics of the plasma membrane during mitosis, affecting the shape and size of mitotic cells and resulting in defects in mitosis progression and generating daughter cells with aberrant cytoarchitecture. In these conditions, F-actin-astral microtubule crosstalk is impaired leading to mitotic spindle misassembly and misorientation, which in turn contributes to chromosome mis-segregation. Mechanistically, we identify the S100 Ca^2+^-binding protein A11 (S100A11) as a key membrane-associated regulator that forms a complex with E-cadherin and LGN to coordinate plasma membrane remodelling with E-cadherin-mediated cell adhesion and LGN-dependent mitotic spindle machinery. We conclude that plasma membrane-mediated maintenance of mammalian epithelial cell identity is crucial for correct execution of polarised cell divisions, genome maintenance and safeguarding tissue integrity.

## Introduction

Cell division requires equal partitioning of DNA into two daughter cells. To achieve this, each dividing cell builds a mitotic spindle that results from a spectacular rearrangement of microtubules allowing proper alignment of chromosomes during metaphase and their equal separation during anaphase^1^. In polarised epithelia of multicellular organisms, the orientation of a correctly assembled mitotic spindle defines the position and fate of daughter cells, representing a fundamental mechanism that ensures the formation of structured and functional tissues^2^. Mitotic spindle orientation is regulated by an evolutionarily conserved ternary protein complex including the G_αi_ subunit of heterotrimeric G proteins, the Leucine-Glycine-Asparagine repeat protein (LGN) and the nuclear mitotic apparatus protein (NuMA)^3^. During mitosis, the G_αi_-LGN-NuMA complex localises at the cell cortex, which facilitates the interaction of NuMA on astral microtubules and with the microtubule-associated minus-end motor Dynein, generating pulling forces on astral microtubules that ensure correct positioning of the mitotic spindle^3–6^. Cumulative evidence shows that the crosstalk between astral microtubules and cortical F-actin is a key mechanism for balancing the cortical forces that ensure correct mitotic spindle and chromosome dynamics, as well as mitosis progression^7–10^. In mammalian epithelia, polarity proteins including E-cadherin, Afadin, ABL1, SAPCD2 and Annexin A1 (ANXA1) have been shown to act as molecular landmarks instructing the polarised patterning of LGN and NuMA at the cell cortex to regulate the balance between planar and perpendicular mitotic spindle orientation^4, 9, 11–13^. Yet, the mechanisms coordinating the interplay between polarity cues, the cell cortex and the mitotic machinery in mammalian epithelial cells remain largely unknown.

During mitosis, cells transduce external cues to reorganise F-actin and microtubules at the cell cortex to control cell shape and balance intracellular tensions, thereby creating an optimal space for mitotic spindle formation and facilitate the subsequent correct alignment of the metaphase chromosomes^14–20^. While most of our knowledge of the mechanics of mitosis in mammalian cells comes from studies in non-polarised cells or cells grown in isolation on adhesive micropatterns^16, 17, 21^, increasing evidence shows that mitotic epithelial cells must maintain their native polarity and geometry to ensure correct mitotic spindle dynamics and chromosome segregation fidelity^22–24^. Mitotic epithelial cells round up to push against tissue confinement ensuring cell mechanics that limits spindle assembly and chromosome segregation errors^25^, while sustaining adherens junctions with their neighbouring cells to maintain epithelial integrity^12, 26–28^. Reciprocally, cell rounding is influenced by tensions and mechanical forces emanating from neighbouring cells, tissue topology as well the growing and remodelling tissue itself^16, 17^. Initial experiments in the amphibian eggs showed that the mitotic spindle aligns along the longest axis of the cell, following Hertwig’s rule^29^. Subsequent in-depth studies have shown that epithelial cell anisotropy determines mitotic spindle orientation, where moderately anisotropic cells follow partially Hertwig’s rule, whereas elongated cells favour division along the major axis^24^. A few planar polarity cues such as Dishevelled and Vangl2 direct mitotic spindle orientation following Hertwig’s rule^30, 31^. Further studies in the *Drosophila* pupal notum showed that LGN and NuMA localise to tricellular junctions to act as cell shape sensors and direct planar cell division along the major axis of the cell^32^. Consistent with this, studies in the *Xenopus* epithelium reported that the mitotic spindle aligns to an axis of cell shape defined by the position of tricellular junctions, which requires functional cell-cell adhesion E-cadherin protein and the localization of LGN to tricellular junctions^33^. However, studies in *canine* kidney MDCK cells showed that E-cadherin and cortical LGN align epithelial cell divisions with tissue tension independently of cell shape, and that the localisation of E-cadherin at the plasma membrane is key to pattern LGN at the cell cortex^12, 27^. The precise molecular mechanisms allowing E-cadherin-mediated adhesion to transduce external cues to ensure correct execution of polarised cell divisions remains not well understood in mammalian epithelial cells.

Mitotic cells undergo dynamic changes in their volume and surface topology, driven by remodelling in the plasma membrane, which responds to mechanical stresses and connects with the cortex to fine-tune intracellular tensions that control the orientation, progression, and outcome of cell division^21, 34^. Depletion of plasma membrane proteins such as the G protein-coupled receptors can inhibit cell division^35, 36^. Other studies reported differences in the cell surface proteome between interphase and metaphase cells^37^. In single HeLa cells, elongation of the plasma membrane is coordinated with cortical localisation of Dynein to centre the mitotic spindle in anaphase and achieve symmetric cell division^38^. In these cells, polar plasma membrane blebbing, stabilises cell shape by relieving actomyosin cortical tensions ensuring the stability of cleavage furrow positioning for correct progression of cytokinesis^39^. In polarised epithelial cells, on the other hand, decrease in intracellular tensions relies on traction forces from neighbouring cells in addition to those emanating from the extracellular matrix^40, 41^, further highlighting the functional requirement of cell-cell signalling for correct execution of polarised cell divisions. However, it remains unknown how mitosis dynamic progression and outcome are coordinated with plasma membrane remodelling and cell-cell adhesion.

Here we exploit a simple monolayer culture system combined with fluorescence time-lapse and confocal imaging to experimentally manipulate the density of mammary epithelial cells and examine how perturbation of cell-cell adhesion influences the orientation, mechanics, and outcome of cell division. We show that cells grown at low density lose their polarised epithelial identity and display aberrant dynamics of the plasma membrane during mitosis, which affects the size and shape of mitotic cells resulting in daughter cells with cytoarchitectural defects. In these non-polarised conditions, cortical actin organisation and its interaction with astral microtubules are impaired. Consequently, cells fail to correctly align the mitotic spindle and chromosomes, leading to delayed mitosis progression and cytokinesis defects. In this experimental context, we investigated the function of the membrane associated protein S100A11 (S100 Ca^2+^-binding protein A11), which we identified recently in a proteomic screening of mitotic mammary epithelial cells^9^. We demonstrate that S100A11 is required for proper plasma membrane remodelling to ensure faithful segregation of chromosomes and generation of equal-sized daughter cells. Mechanistically, we show that S100A11 forms a complex with E-cadherin and LGN to instruct correct E-cadherin-mediated adhesion and lateral patterning of the LGN-mediated spindle orientation machinery, thereby ensuring planar cell division. S100A11 depletion phenocopies the mitotic defects observed in non-polarised cells and alters epithelial integrity. Collectively, the present experiments shed new light onto the importance of epithelial identity maintenance for correct dynamics, mechanics, and outcome of polarised cell divisions, and establish S100A11-mediated plasma membrane remodelling as a mechanism coordinating the mechanochemical crosstalk between cell-cell adhesion, the cell cortex and mitotic spindle machinery in mammalian epithelial cells.

## Results

### Epithelial cell density-dependent plasma membrane remodelling influences mitosis progression and outcome

To examine the functional requirement of cell-cell adhesion for correct plasma membrane remodelling during cell division, we used human MCF-10A mammary epithelial cells cultured at optimal or low density (Figure 1A). Characterisation of MCF-10A monolayers 72 hours (h) after plating reveals that cells cultured at optimal density establish E-cadherin adherens junctions, whereas cells cultured at low density lose their polarised epithelial identity and fail to accumulate E-cadherin at the cell surface both in interphase and metaphase, allowing us to define optimal-density MCF-10A cells as polarised, and low-density MCF-10A cells as non-polarised (Figures 1A and 1B). Next, we performed live imaging in polarised and non-polarised MCF-10A cells, in which we labelled the plasma membrane with CellMask^TM^ and DNA with Hoechst 33342. Consistent with previous studies ^42^, CellMask^TM^ displays a homogeneous, circumferential distribution at the cell surface of polarised mitotic cells, which generate equal-sized daughter cells at cytokinesis (Figures 1C-1E; Movie S1). In non-polarised mitotic cells, however, the plasma membrane has aberrant dynamics with ∼52% of cells displaying a unilateral accumulation of CellMask^TM^ at the cell surface (Figures 1C-1E; Movie S2; Movie S3). Loss of cell-cell adhesion integrity affects cell size and shape and leads to polar, asymmetric elongation of the plasma membrane during anaphase in ∼77% of mitotic cells (Figures 1C and 1F). In these conditions, sister chromatids are off-centred from anaphase until telophase, which affects the position of the cleavage furrow and generates unequal-sized daughter cells at cytokinesis in ∼44% of non-polarised mitotic cells (Figures 1G and 1H), consistent with previous findings in HeLa cells^38^. During anaphase-to-telophase transition, the plasma membrane of non-polarised cells forms blebs in the polar area opposing the expanding cell cortex, but this fails to re-centre the sister chromatids or stabilise the cleavage furrow (Figure 1C), which disagrees with previous studies showing that polar membrane blebbing influences the position and stability of the mitotic spindle and cleavage furrow^38, 39^. Interestingly, a close examination of the CellMask^TM^ labelling in non-polarised cells reveals an accumulation of cytoplasmic vesicles, some of which localise to the cleavage furrow (Figure 1C). Thus, asymmetric membrane elongation may involve both remodelling and *de novo* incorporation of membrane components into the cleavage furrow. We also observed significant defects in the dynamics of cell division in a vast majority of non-polarised cells as compared to polarised cells, with the proportion of cells that completed mitosis decreases significantly in low-density cell cultures (non-polarised: ∼50% versus polarised: ∼94%) (Figure 1I). In non-polarised cells that completed their division, the duration of mitosis is extended (Figure 1C), as revealed by increased the average transition time from nuclear envelope breakdown (NEBD) to cytokinesis (non-polarised: ∼98 min versus polarised: ∼62 min) (Figure 1J), consistent with a recent study in MDCK cells showing that epithelial density influences progression of the cell cycle^43^. Together, these data indicate that cell-cell adhesion integrity and plasma membrane remodelling are coordinated to maintain epithelial identity and ensure correct mitotic mechanics and progression as well as symmetric cytokinesis.

**Figure 1.**
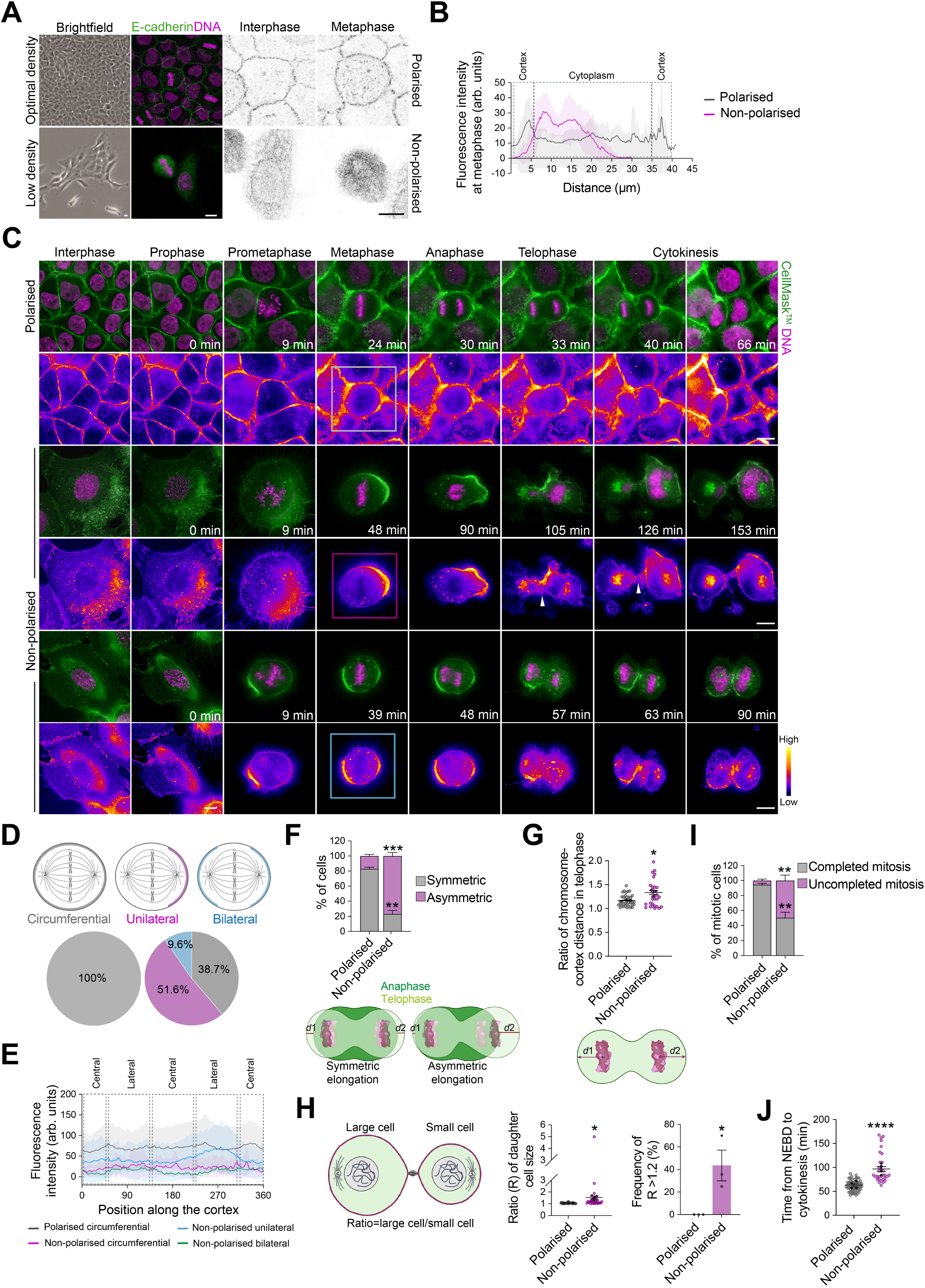
Perturbation of cell-cell adhesion integrity results in asymmetric plasma membrane elongation and defects in mitosis progression and outcome. (**A**) Confocal images of representative polarised and non-polarised MCF-10A cells stained for E-cadherin (green) and counterstained with Hoechst 33342 (DNA, magenta). (**B**) Average cortical and cytoplasmic fluorescence intensity profiles of E-cadherin in polarised and non-polarised metaphase cells (polarised: *n* = 26 cells; non-polarised: *n* = 24 cells). (**C**) Time-lapse images of representative polarised and non-polarised cells. The plasma membrane is labelled with CellMask^TM^ (green), and DNA with Hoechst 33342 (magenta), 10 min and 30 min before acquisition, respectively. Arrowheads indicate accumulation of vesicles at the cleavage furrow in non-polarised cells. Insets indicate metaphase cells showing the localisation phenotypes of CellMask^TM^ quantified in D: Circumferential (grey); Unilateral (pink); Bilateral (blue). (**D**) Percentage of CellMask^TM^ labelling distribution at the cell surface during metaphase in polarised and non-polarised cells (polarised: *n* = 30 cells; non-polarised: *n* = 30 cells). (**E**) Average cortical fluorescence intensity profiles of CellMask^TM^ from polarised and non-polarised cells (polarised: *n* = 30 cells; non-polarised: *n* = 30 cells). (**F**) Percentage of cells with symmetric and asymmetric plasma membrane elongation, (polarised: *n* = 34 cells; non-polarised: *n* = 32 cells). Distance (*d*) was measured as described on the illustration. Two-sided *t*-test, symmetric: \*\*\**P* = 0.0003; asymmetric: \*\**P* = 0.0012. (**G**) Ratio of chromosome to cortex distance (*d*) in telophase cells (polarised: *n* = 37 cells; non-polarised: *n* = 30 cells). Two-sided t-test, \**P* = 0.014. *d* was measured as described on the illustration. (**H**) Left: ratio (R) of daughter cell size. *t*-test, \**P* = 0.018. Right: frequency of R < 1.2. Two-sided *t*-test, \**P* = 0.0193. (polarised: *n* = 35 cells; non-polarised: *n* = 30 cells). Ratio was measured as described on the illustration. (**I**) Percentage of cells with complete and incomplete mitosis (polarised: *n* = 63 cells; non-polarised: *n* = 67 cells). Two-sided *t*-test, \*\**P* = 0.0013. (**J**) Time from nuclear envelope breakdown (NEBD) to cytokinesis (polarised: *n* = 58 cells; non-polarised: *n* = 34 cells). Two-sided *t*-test, \*\*\*\**P* < 0.0001. All data are presented as mean ± s.e.m. from 3 or 4 independent experiments. arb. units (arbitrary units). All scale bars, 10 µm. Source data are provided as a Source Data file.

### Epithelial cell density-dependent mitotic spindle dynamics and chromosome segregation fidelity

The results above strongly suggest that epithelial cell density influences the assembly and alignment of the mitotic spindle, which in turn ensures correct chromosome dynamics and segregation. To test this hypothesis, we performed live imaging of polarised and non-polarised MCF-10A cells, in which we labelled microtubules with SiR-tubulin^44^ and DNA with Hoechst 33342 (Figure 2A). We identified several mitotic progression defects in non-polarised cells (Figures 2A and 2B; Movie S4; Movie S5; Movie S6; Movie S7). We observed significant defects in chromosome alignment and segregation in ∼60% and ∼27% of non-polarised cells, respectively, leading to a significant increase in the proportion of daughter cells inheriting micronuclei at cytokinesis (non-polarised: ∼15 % versus polarised: 0%) (Figures 2A and 2B, Movie S5). The morphology of the mitotic spindle is also affected, with ∼35% of non-polarised cells displaying abnormal bipolar spindles (Figures 2A and 2B; Movie S7). During metaphase, polarised cells align their mitotic spindle parallel to the substratum plane (∼4°), whereas non-polarised cells display spindle orientation defects (∼10°), which persist during anaphase (non-polarised: ∼9° versus polarised: ∼4°) (Figure 2C). We further confirmed these observations using immunofluorescence and confocal imaging (Figure 2D). Our live imaging experiments also show that ∼93% of non-polarised cells display excessive oscillations of the mitotic spindle relative to the *z* axis between successive time frames during metaphase, whereas in polarised cells the spindle does not display notable oscillatory *z* rotations and remains in a planar position (Figures 2E-2G). There is also an increase in the oscillatory rotations of the mitotic spindle in the *xy* plane in non-polarise cells as compared to polarised cells, but the difference is not significant (Figures 2H-2J). Nonetheless, while all investigated polarised cells align their mitotic spindle following Hertwig’s rule in the *xy* plane, only ∼28% of non-polarised cells follow this rule (Figures S1A and S1B). Thus, cell-cell adhesion integrity ensures correct mitotic spindle assembly and dynamic orientation, and in turn faithful chromosome segregation in dividing mammary epithelial cells.

**Figure 2.**
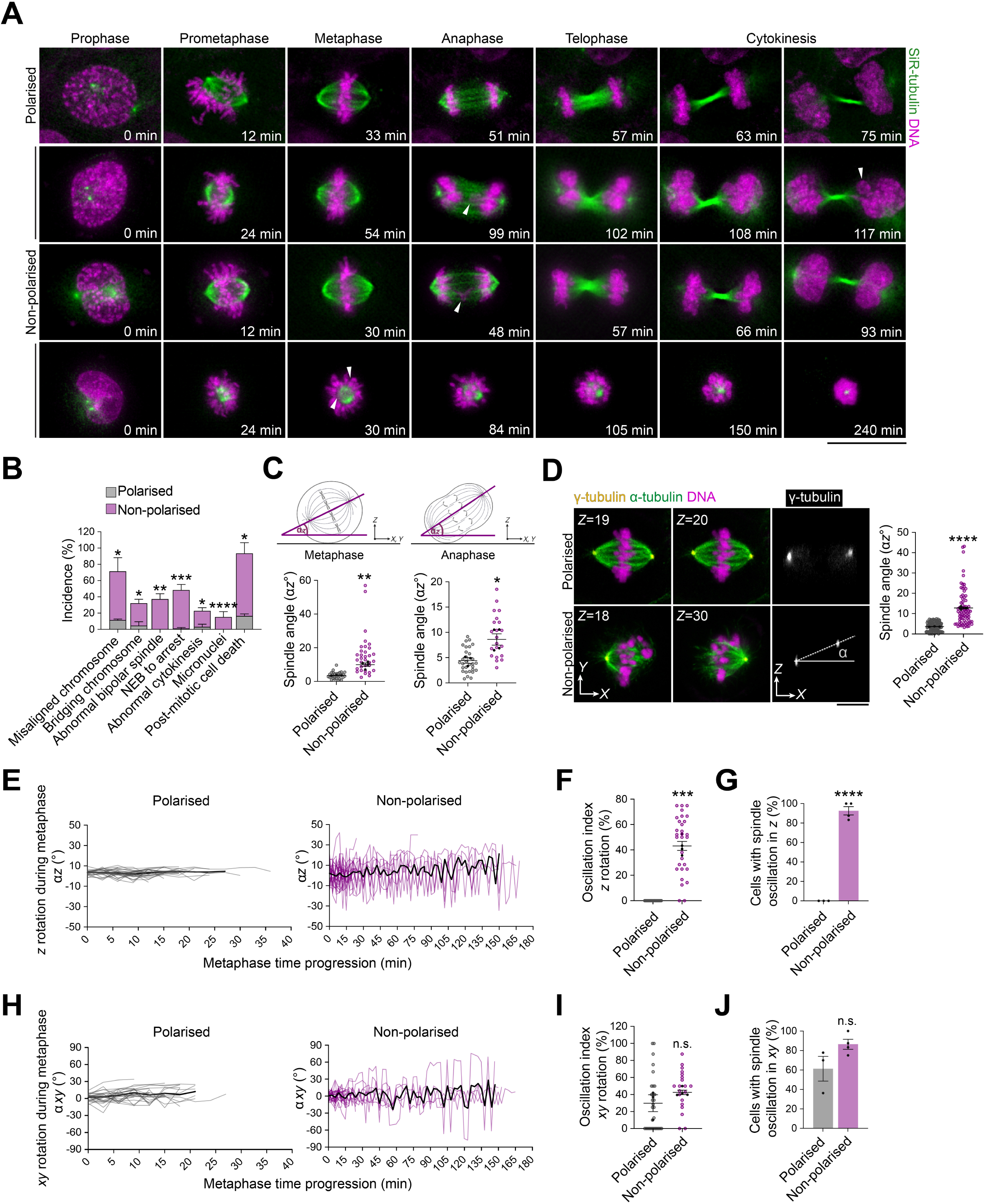
Perturbation of cell-cell adhesion integrity impairs mitotic spindle dynamics and chromosome segregation. (**A**) Time-lapse images of representative polarised and non-polarised MCF-10A cells. Microtubules are labelled with SiR-tubulin (green) and DNA Hoechst 33342 (magenta), 3 h and 30 min before acquisition, respectively. Arrowheads indicate the phenotypes listed in B. Scale bar, 10 µm. **(B)** Percentage of cells with defects in mitotic spindle assembly, chromosome alignment and segregation, and mitotic progression and outcome (polarised: *n* = 61 cells; non-polarised: *n* = 73 cells). Two-sided *t*-test, misaligned chromosome: \**P* = 0.043; bridging chromosome: \**P* = 0.029; abnormal bipolar spindle: \*\**P* = 0.0087; NEB-to-arrest: \*\*\**P* = 0.0006; abnormal cytokinesis: \**P* = 0.0298; micronuclei: \*\*\*\**P* <0.0001; and post-mitotic cell death: \**P* = 0.0462. **(C)** Mitotic spindle angles α*z* from time-lapse images in polarised and non-polarised cells at metaphase (left) (polarised: *n* = 30 cells; non-polarised: *n* = 30 cells); and anaphase (right) (polarised: *n* = 30 cells; non-polarised: *n* = 21 cells). Two-sided *t*-test, metaphase: \*\**P* = 0.0034; anaphase \**P* = 0.0316. **(D)** Left: confocal images of representative polarised and non-polarised cells stained for α-tubulin (green) and anti-γ-tubulin (yellow), and counterstained with Hoechst 33342 (DNA, magenta). Scale bar, 10 µm. Right: mitotic spindle angle α*z* in fixed polarised and non-polarised cells during metaphase (polarised: *n* = 103 cells; non-polarised: *n* = 74 cells). Two-sided *t*-test, \*\*\*\**P* < 0.0001. **(E)** Dynamics of z orientation (α*z*) in polarised cells (left) and non-polarised cells (right), (polarised: *n* = 30 cells; non-polarised: *n* = 30 cells). Black lines represent the average spindle angles. **(F)** Oscillation index in polarised and non-polarised cells, in the *z* axis (polarised: *n* = 30 cells; non-polarised: *n* = 30 cells). Two-sided *t*-test, \*\*\**P* = 0.0002. **(G)** Percentage of cells with spindle oscillation in the z axis (polarised: *n* = 30 cells; non-polarised: *n* = 30 cells). Two-sided *t*-test, \*\*\*\**P* < 0.0001. **(H)** Dynamics of *xy* orientation (α*xy*) in polarised cells (left) and non-polarised cells (right), (polarised: *n* = 30 cells; non-polarised: *n* = 28 cells). Black lines represent the average spindle angles. **(I)** Oscillation index in polarised and non-polarised cells, in the *xy* axis (polarised: *n* = 30 cells; non-polarised: *n* = 30 cells). Two-sided *t*-test, *P* = 0.203. **(J)** Percentage of cells with spindle oscillation in the *xy* axis (polarised: *n* = 30 cells; non-polarised: *n* = 29 cells). Two-sided *t*-test, *P* = 0.0968. All data are presented as mean ± s.e.m. from 3 or 4 independent experiments. n.s. (not significant). Source data are provided as a Source Data file.

### Epithelial cell density-dependent re-organisation of cortical actin during mitosis

Dynamic re-organisation of actin cytoskeleton into a uniform, contractile cortex at the cell surface at mitotic entry ensures proper assembly and orientation of the mitotic spindle by polarising cortical force-generating proteins that pulls on astral microtubules^4, 9, 10^. This cortical actin network is important for accurate control of cell rounding, which peaks during metaphase^16, 17^. In epithelial cells, mitotic rounding defects result in abnormal spindle assembly and orientation, division asymmetries as well as delayed mitotic progression^14, 45^. To investigate how perturbation of cell-cell adhesion affects cortical actin re-organisation during mitosis, we performed live imaging in MCF-10A cells stably expressing Lifeact-mCherry and labelled with SiR-tubulin (Figure 3A; Movie S8; Movie S9). We measured cortical thickness in metaphase, which is a key readout of cortical actin architecture^15^, and found that perturbation of cell-cell adhesion integrity leads to an increase in cortex thickness (Figure 3B). F-actin fluorescence intensity also increases in both the cortical and subcortical regions of non-polarised metaphase cells (Figure 3C). Consequently, non-polarised cells are rounder, and the rounding-up process is faster as compared to polarised cells (non-polarised: ∼20 min versus polarised: ∼28 min) (Figures 3D and 3E). We confirmed these observations by immunofluorescence and confocal imaging, which also reveal a decrease in the depth of the subcortical actin cloud in non-polarised cells (non-polarised: ∼5 μm versus polarised: ∼8 μm) (Figures 3F and 3G; Figures S2A-S2C). This is associated with impaired astral microtubule length in non-polarised cells (non-polarised: ∼19 nm versus polarised: ∼16 nm) while their relative intensity remains unaffected, suggesting their stabilisation (Figures 3H and 3I). Finally, the length of the mitotic spindle as well as the fluorescence intensity of spindle microtubules increase in non-polarised cells as compared to polarised cells (Figures S2D and S2E). Thus, cell-cell adhesion defines the dynamic re-organisation of the actin cortex and its interaction with astral microtubules to ensure correct mitotic spindle dynamics and mitotic cell rounding.

**Figure 3.**
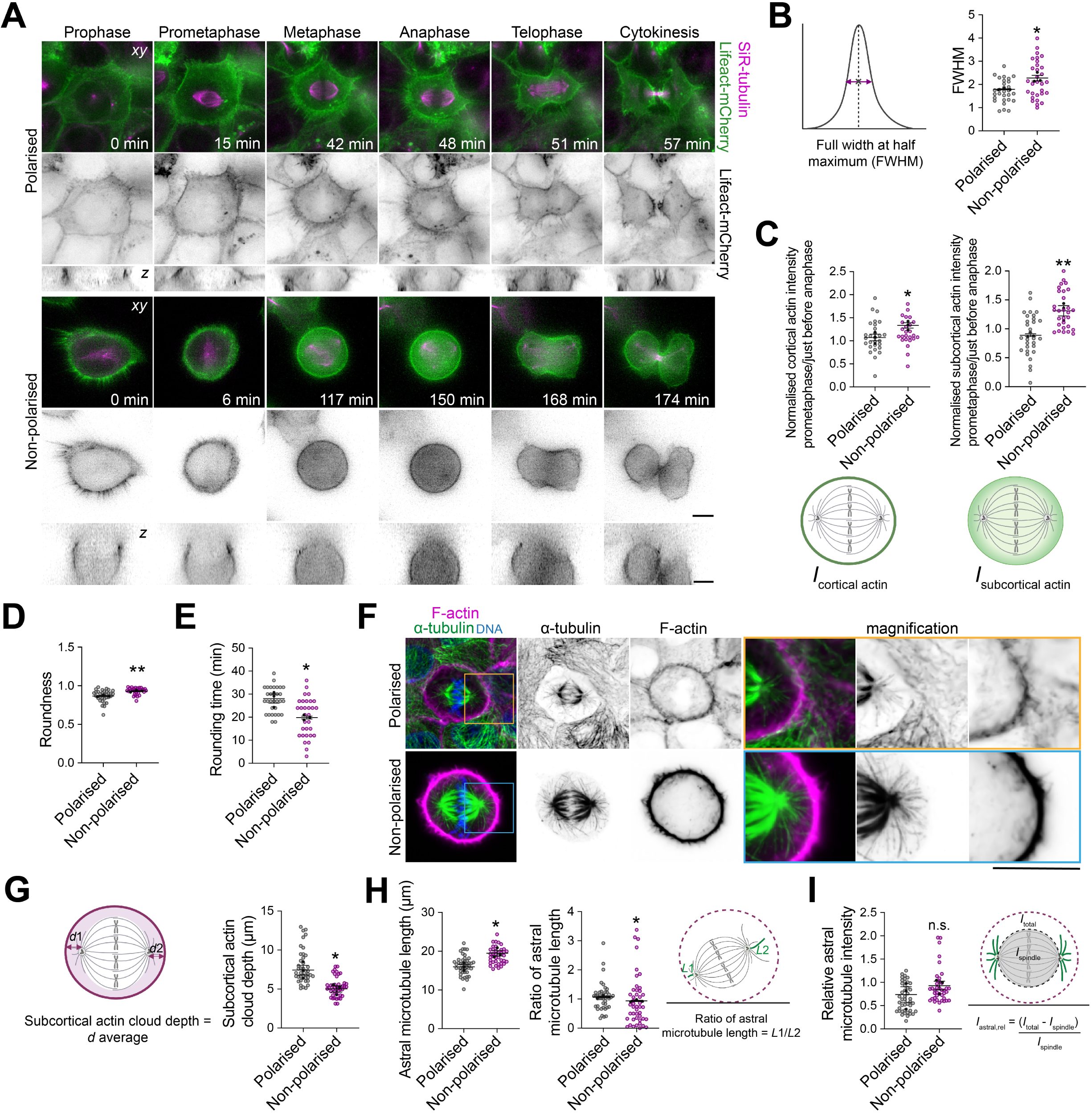
Perturbation of cell-cell adhesion integrity impairs F-actin and astral microtubule organisation and crosstalk. (**A**) Time-lapse images of representative clonal polarised and non-polarised cells MCF-10A cells stably expressing Lifeact-mCherry (green). Microtubules labelled with SiR-tubulin (magenta) 3 h before imaging. Scale bars, 10 µm. **(B)** Full width at half maximum (FWHM) for measurement of actin stiffness (polarised: *n* = 29 cells; non-polarised: *n* = 30 cells). Two-sided *t*-test, \**P* = 0.0169. **(C)** Ratio of fluorescence intensity of cortical (left) and subcortical (right) actin changes between prometaphase and last frame of metaphase: cortical (polarised: *n* = 30 cells; non-polarised: *n* = 29 cells); subcortical (polarised: *n* = 28 cells; non-polarised: *n* = 25 cells). Two-sided *t*-test, cortical \**P* = 0.044; subcortical \*\**P* = 0.010. **(D)** Roundness of metaphase cells (polarised: *n* = 33 cells; non-polarised: *n* = 29 cells). Two-sided *t*-test, \*\**P* = 0.0035. **(E)** Rounding time for polarised and non-polarised cells (polarised: *n* = 31 cells; non-polarised: *n* = 31 cells). Two-sided *t*-test, \**P* = 0.0159. **(F)** Confocal images of representative polarised and non-polarised cells stained for F-actin (magenta) and α-Tubulin (green); and counterstained with Hoechst 33342 (DNA, blue). Scale bar, 10 µm. **(G)** Average subcortical actin cloud depth (*d*) in polarised and non-polarised cells (polarised: *n* = 41 cells; non-polarised: *n* = 43 cells). Two-sided *t*-test, \**P* = 0.029. *d*1 and *d*2 were measured as described on the illustration. **(H)** Left: Astral microtubule length (polarised: *n* = 42 cells; non-polarised: *n* = 39 cells); right: ratio of astral microtubule length (*L*) (polarised: *n* = 42 cells; non-polarised: *n* = 45 cells). Two-sided *t*-test, astral microtubule length: \**P* = 0172; ratio of astral microtubule length: \**P* = 0176. *L*1 and *L*2 were measured as described on the illustration. **(I)** Relative (rel) fluorescence intensities of astral microtubules (*I*_astral, rel_) in polarised and non-polarised cells (polarised: *n* = 43 cells; non-polarised: *n* = 39 cells). Two-sided *t*-test, *P* = 0.37. *I*_total_ and *I*_spindle_ were measured as described on the illustration. All data are presented as mean ± s.e.m. from 3 or 4 independent experiments. n.s. (not significant). Source data are provided as a Source Data file.

### S100A11 coordinates plasma membrane remodelling with cell-cell adhesion integrity and cortical cytoskeleton organisation during mitosis

We recently characterised the LGN cortical interactome in mitotic mammary epithelial cells, where we identified ANXA1 as a polarity cue that controls planar mitotic spindle orientation^9^. Our proteomic screening also identified S100A11, an established partner of ANXA1^46^. S100A11 associates to the plasma membrane in a Ca^2+^-dependent manner and has been shown to regulate plasma membrane repair and integrity during cell migration, by regulating F-actin dynamics and plasma membrane remodelling in migrating breast cancer cells^47^. S100A11 is enriched in pseudopodia of metastatic cancer cells and is required for the formation of actin-dependent pseudopodia protrusions and tumour cell migration^48^. Furthermore, S100A11 has been identified as an interactor of E-cadherin at adherens junctions^49^. These findings prompted us to test whether S100A11 function in the regulation of plasma membrane remodelling may influence cell-cell adhesion integrity and F-actin re-organisation during mitosis. We first performed live imaging to characterise the dynamic localisation of S100A11 in MCF-10A cells stably expressing GFP-S100A11. We show that GFP-S100A11 distributes in the cytoplasm and accumulates at the cell surface throughout the cell cycle (Figure 4A; Movie S10). We confirmed these observations by immunofluorescence and confocal imaging (Figure S3A). To examine the role of S100A11 in plasma membrane remodelling during mitosis, we performed live imaging in control and S100A11-depleted MCF-10A cells grown at optimal density, which we labelled with CellMask^TM^ (Figure 4B; Movie S11; Movie S12). We observed unequal distribution of CellMask^TM^ in a vast majority of S100A11-depleted cells (si-S100A11#1: ∼79% versus si-Control: ∼26%), indicating defects in plasma membrane remodelling (Figures 4B and 4C). The proportion of cells that completed mitosis decreases significantly upon S100A11 knockdown (si-S100A11#1: ∼61% versus si-Control: ∼98%) (Figure 4D). S100A11-depleted cells that completed mitosis display chromosome alignment and segregation defects in ∼16% and ∼67% of cells, respectively (Figure 4E). Immunofluorescence and confocal imaging reveal that S100A11 knockdown impairs the localisation of E-cadherin at adherens junctions at 48 h post-S100A11 knockdown, while the total cellular levels of E-cadherin do not change (Figures 4F and 4G). These defects in cell-cell adhesion persist at 72 h post-S100A11 depletion (Figure S3B). A close examination of the localisation of E-cadherin in metaphase cells shows that the protein translocates to the cytoplasm and fails to cluster properly at adherens junctions of S100A11-depleted cells (Figures 4H and 4I). This impairs cell-cell adhesion integrity and results in significant defects in the cytoarchitecture of S100A11-depeleted cells (Figures 4J and 4K). Consistent with this, S100A11 knockdown abrogates the formation of single tight F-actin bundles at cell-cell contacts (Figures S3C and S3D) and affects the organisation of cortical F-actin and its interaction with astral microtubules during metaphase, of which the length is also affected (Figures S3E-3G). Our results indicate that S100A11 is a key membrane-associated protein that coordinates the crosstalk between the plasma membrane, cell-cell adhesion, and the cell cortex during mitosis.

**Figure 4.**
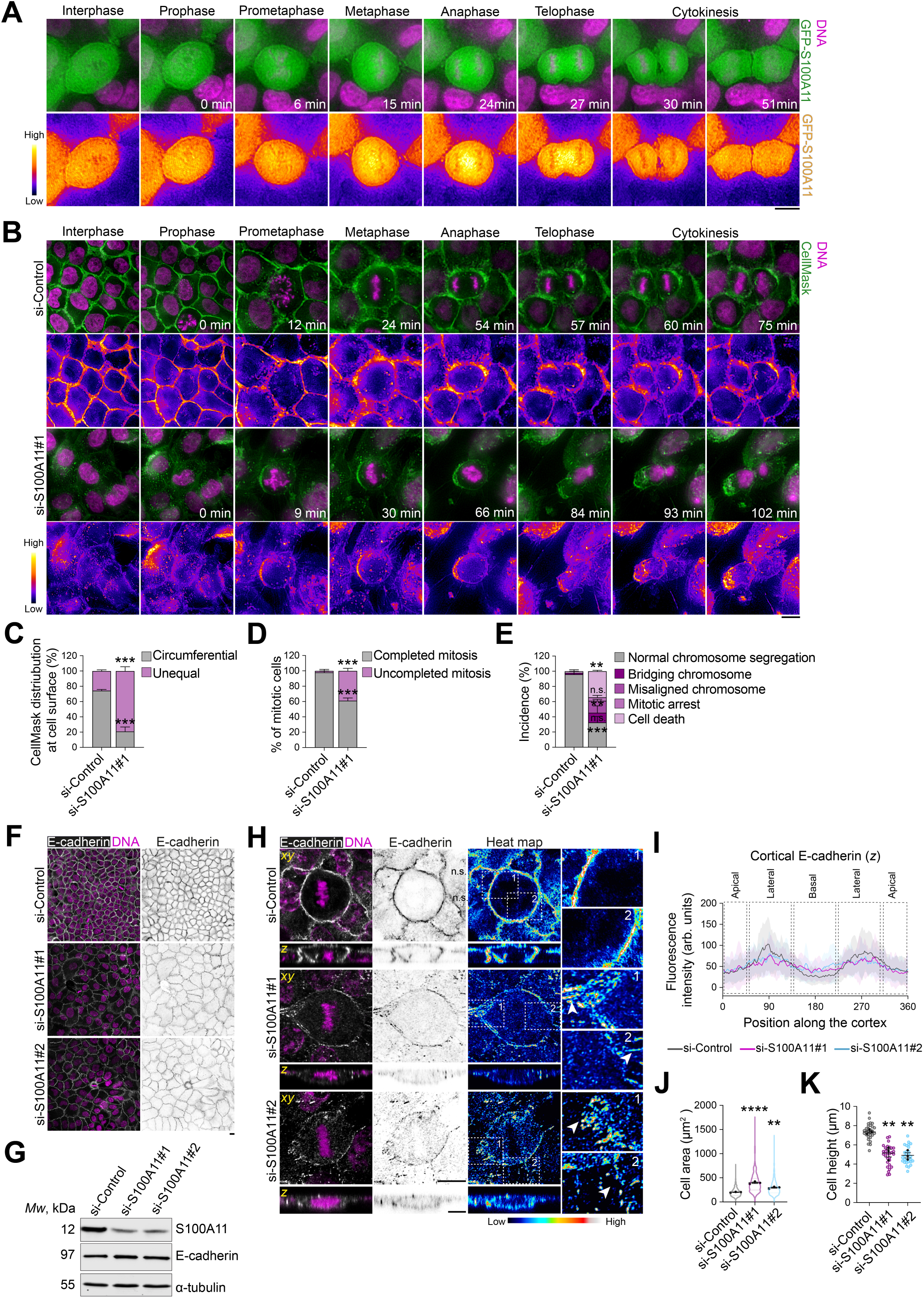
S100A11 is required for correct plasma membrane remodelling and cell-cell adhesion integrity during mitosis. (**A**) Time-lapse images of representative clonal MCF-10A cells stably expressing GFP-S100A11 (green). DNA is labelled with Hoechst 33342 (magenta). Scale bar, 10 µm. **(B**) Time-lapse images of representative si-Control-or si-S100A11#1-treated MCF-10A cells, labelled with CellMask^TM^ (plasma membrane, green) and Hoechst 33342 (DNA, Magenta), 10 min and 30 min before acquisition, respectively. Scale bar, 10 µm. **(C)** Percentage of CellMask^TM^ labelling distribution at the cell surface during metaphase in siRNA-transfected cells (si-Control: *n* = 28 cells; si-S100A11#1: *n* = 59 cells). Two-sided *t*-test, circumferential: \*\*\**P* = 0.0008; unequal: \*\*\**P* = 0.0008. **(D)** Incidence of completed and uncompleted mitosis in siRNA-transfected cells (si-Control: *n* = 28 cells; si-S100A11#1: *n* = 59 cells). Two-sided *t*-test, completed: \*\*\**P* = 0.0007; uncompleted: \*\*\**P* = 0.0007. **(E)** Percentage of mitotic defects in siRNA-transfected cells (si-Control: *n* = 28 cells; si-S100A11#1: *n* = 59 cells). Two-sided *t*-test, normal chromosome segregation: \*\*\**P* = 0.0002; misaligned chromosome: \*\**P* = 0.0012; bridging chromosome: *P* = 0.308; mitotic arrest: *P* = 0.172; cell death: \*\**P* = 0.002. **(F)** Confocal images of representative si-Control-, si-S100A11#1, si-S100A11#2-treated cells, stained for E-cadherin (grey) and counterstained with Hoechst 33342 (DNA, magenta). Scale bar, 10 µm. **(G)** Western blotting of extracts from siRNA-transfected cells. Blots are stained for S100A11 and E-cadherin, and α-tubulin as a loading control. **(H)** Confocal images of representative si-Control-, si-S100A11#1, si-S100A11#2-treated metaphase cells stained for E-cadherin (grey) and counterstained with Hoechst 33342 (DNA, magenta). Scale bars, 10 µm. **(l)** Average cortical fluorescence intensity profiles of E-cadherin in siRNA-transfected metaphase cells (si-Control: n = 30; si-S100A11#1: n = 30; si-S100A11#2: n = 27). **(J)** Cell area in siRNA-transfected cells (si-Control: *n* = 1699 cells; si-S100A11#1: *n* = 634 cells; si-S100A11#2: *n* = 1110 cells). One-way ANOVA with Dunnett’s test, \*\*\*\**P* < 0.0001, \*\**P* = 0.002. **(K)** Cell height in siRNA-transfected cells (si-Control: *n* = 30 cells; si-S100A11#1: *n* = 30 cells; si-S100A11#2: *n* = 29 cells). One-way ANOVA with Dunnett’s test, \*\**P* = 0.0032, \*\**P* = 0.0019. All data are presented as mean ± s.e.m. from 3 or 4 independent experiments. n.s. (not significant). Source data are provided as a Source Data file.

### S100A11 forms a complex with E-cadherin and LGN to control mitotic spindle orientation

Both cortical F-actin and E-cadherin interact with LGN to regulate the localisation and function of NuMA-Dynein-Dynactin to generate cortical forces that orient the mitotic spindle in epithelial cells^4, 12^. Our results described above and previous findings of other labs showing that S100A11 interacts with E-cadherin^49^ and F-actin^48^, prompted us to test whether S100A11 regulates LGN-mediated mitotic spindle orientation. First, we performed affinity purification to isolate the S100A11 complex from stable MCF-10A cells expressing GFP-S100A11 and arrested in metaphase (Figure 5A). We validated synchronisation efficiency by accessing the accumulation of phospho-Histone H3 (Figure S4). Our pull-down assays combined with western blotting analysis reveal that GFP-S100A11 co-precipitates LGN and E-cadherin (Figure 5A). Previously, endogenous S100A11 has been co-purified in a reciprocal manner with GFP-LGN^9^ and E-cadherin-BirA*^49^. Furthermore, we reveal that endogenous S100A11 co-localises with E-cadherin to the cell surface in polarised epithelial cells, whereas the two proteins co-translocate to the cytoplasm with E-cadherin when cell-cell adhesions are disrupted (Figures 5B and 5C). Next, we performed immunofluorescence and confocal imaging to evaluate the extent to which S100A11 affects the localisation of LGN at the lateral cortex during metaphase. S100A11 knockdown impairs the bilateral cortical distribution of LGN observed in control cells, with a significant increase in the proportion of S100A11-depleted cells displaying a unilateral accumulation of LGN (si-S100A11#1: ∼45%; si-S100A11#2: ∼52%; si-Control: ∼11%) (Figures 5D and 5E). These results further suggest that S100A11 regulates mitotic spindle orientation. We tested this hypothesis directly by immunofluorescence and confocal imaging and found that S100A11 knockdown results in mitotic spindle misorientation, with spindle angles α*z* increasing from ∼4° in controls to ∼10° and ∼14° in cells treated with si-S100A11#1 and si-S100A11#2, respectively (Figures 5F and 5G). Interestingly, E-cadherin knockdown impairs the recruitment of LGN to the cell cortex (Figures 5H-5J), but moderately affects S100A11 localisation at the plasma membrane (Figures 5K and 5L), suggesting that S100A11 may act upstream of E-cadherin to control the mitotic spindle orientation machinery.

**Figure 5.**
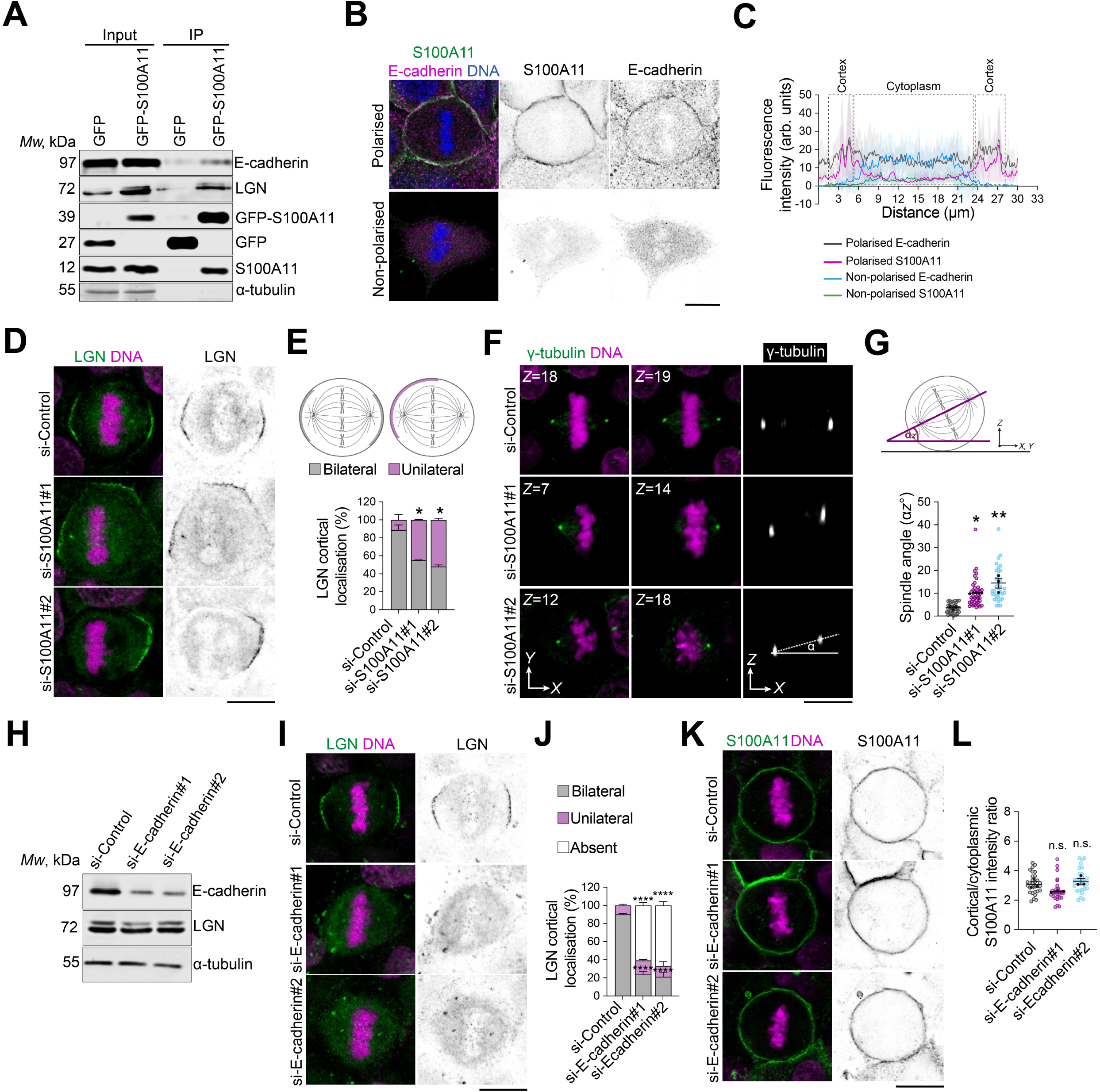
S100A11 complexes with E-cadherin and LGN to regulate E-cadherin-LGN mediated mitotic spindle orientation. (**A**) GFP-S100A11, E-cadherin and LGN co-immunoprecipitates from clonal MCF-10A stably expressing GFP-S100A11, arrested in metaphase. Lysates were subjected to affinity purification with GFP-Trap beads. The immunoprecipitates (IP) were analysed by western blotting. **(B)** Confocal images of representative polarised and non-polarised MCF-10A cells stained for S100A11 (green) and E-cadherin (magenta); and counterstained with Hoechst 33342 (DNA, blue). Scale bar, 10 µm. **(C)** Average cortical and cytoplasmic relative fluorescence intensity profiles of E-cadherin and S100A11 in 10 representative polarised cells and 10 representative non-olarised cells, during metaphase. **(D)** Confocal images of representative si-Control-, si-S100A11#1-and si-S100A11#2-treated metaphase cells stained for LGN (green) and counterstained with Hoechst 33342 (DNA, magenta). Scale bar, 10 µm. **(E)** Percentage of cortical localisation of LGN in siRNA-transfected cells (si-Control: *n* = 64 cells; si-S100A11#1: *n* = 64 cells; si-S100A11#2: *n* = 49 cells). Two-way ANOVA with Tukey’s test, \**P* = 0.016; \**P* = 0.0104; *P* = 0.924, respectively. **(F)** Confocal images of representative si-Control-, si-S100A11#1, si-S100A11#2-treated metaphase cells stained for γ-tubulin (green) and counterstained with Hoechst 33342 (DNA, magenta). Scale bar, 10 µm. **(G)** Mitotic spindle angle for siRNA-transfected cells during metaphase, (si-Control: *n* = 37 cells; si-S100A11#1: *n* = 41 cells; si-S100A11#2: *n* = 36 cells). One-way ANOVA with Dunnett’s test, \**P* = 0.025; \*\**P* = 0.002, respectively. **(H)** Western blotting of extracts from siRNA-transfected cells. Blots are stained with E-cadherin, LGN and α-tubulin as a loading control. **(I)** Confocal images of representative si-Control-, si-E-cadherin#1, si-E-cadherin#2-treated metaphase cells stained for LGN (green) and counterstained with Hoechst 33342 (DNA, magenta). Scale bars, 10 µm. **(J)** Percentage of cortical localisation of LGN in siRNA-transfected cells (si-Control: *n* = 39 cells; si-E-cadherin#1: *n* = 51 cells; si-E-cadherin #2: *n* = 43 cells). Two-way ANOVA with Dunnett’s test, bilateral: \*\*\*\**P* <0.0001 and \*\*\*\**P* <0.0001; unilateral: *P* = 0.379 and *P* = 0.891; absent: \*\*\*\**P* < 0.0001 and \*\*\*\**P* <0.0001. **(K)** Confocal images of representative si-Control-, si-E-cadherin#1, si-E-cadherin#2-treated metaphase cells stained for S100A11 (green) and counterstained with Hoechst 33342 (DNA, magenta). Scale bars, 10 µm. **(L)** Cortical to cytoplasm ratio of S100A11 fluorescence intensities in siRNA-transfected cells (si-Control: *n* = 30 cells; si-E-cadherin#1: *n* = 30 cells; si-E-cadherin #2: *n* = 30 cells). One-way ANOVA with Dunnett’s test, *P* = 0.114; *P* = 0.589. All data are presented as mean ± s.e.m. from 3 or 4 independent experiments. n.s. (not significant). Source data are provided as a Source Data file.

## Discussion

Polarised cell divisions are crucial for mammalian epithelial tissue differentiation, integrity and function, by defining the fate and position of daughter cells^2^. Recent evidence has shown that epithelial polarity influences the orientation, progression, and outcome of cell division, including how chromosomes segregate to daughter cells^2, 12, 22, 23, 27^. However, most of our mechanistic knowledge of how cell shape dictates the dynamic progression of mammalian cell divisions has advanced, largely from studies in non-polarised or isolated cells grown on adhesive micropatterns^17, 24, 38^. While these studies have extensively described the influence of adhesive substrates and the extracellular matrix on the mechanics of mitosis by regulating the crosstalk between cortical cytoskeleton and the mitotic machinery^16, 17^, how these mechanisms are coordinated with plasma membrane signalling and cell-cell adhesion remains poorly defined. Our present experiments in mammary epithelial cells uncover a functional interplay between plasma membrane remodelling and cell-cell adhesion, which maintains epithelial cell identity to control the dynamics and outcome of polarised cell divisions. We further report that cell-cell adhesion integrity maintains correct plasma membrane remodelling and F-actin-astral microtubule crosstalk to control the shape and size of mitotic cells, which in turn ensures proper mitotic spindle dynamics and faithful chromosome segregation, as well as symmetric separation of daughter cells during cytokinesis. At the molecular level, we identify S100A11 as an upstream membrane-associated cue that coordinates correct plasma membrane remodelling with cell-cell adhesion integrity and cortical cytoskeleton re-organisation during mitosis. We demonstrate that S100A11 complexes with E-cadherin and LGN to direct their polarised localisation to the lateral cortex for correct orientation, progression, and outcome of mammary epithelial cell divisions. Our findings shed new light on the function of the plasma membrane as a molecular signalling platform coordinating correct mechanochemical crosstalk between mitotic epithelial cells and their tissue environment to maintain their epithelial identity and execute error-free cell divisions.

Most studies addressing the role of external cues in the regulation of mammalian cell divisions have been carried out in isolated cells grown on adhesive micropatterns, a method pioneered by Bornens and Théry^50^. These studies have shown that mitotic focal/substrate adhesion, maintained by integrin signalling, is key for correct chromosome and mitotic spindle dynamics and cytokinesis^22, 51–53^. Nonetheless, cell-cell adhesion integrity is essential for epithelial cells to acquire a defined shape and polarity, which are maintained to ensure correct execution of the mitotic machinery^16, 21, 54^. Disruption of tissue architecture in several mammalian epithelial tissues such as skin, liver, prostate, and mammary gland, affects cell shape and polarity leading to defects in chromosome segregation and mitotic dynamics^22, 23^. Our experiments show that perturbation of cell-cell adhesion in mammary epithelial cells cultured at low-density abrogates E-cadherin localisation to the cell surface, which affects cell shape throughout mitosis. Our results suggest, but do not prove, that loss of cell-cell adhesion integrity affects plasma membrane remodelling, which elongates asymmetrically to generate unequal-sized daughter cells at cytokinesis. This is in sharp contrast with studies in isolated HeLa cells, showing that Anillin-RANGTP-dependent plasma membrane asymmetric elongation corrects mitotic spindle positioning defects during late anaphase and ensure symmetric cell division^38^. Our observations that asymmetric elongation of the plasma membrane upon loss of cell-cell adhesion integrity results in mitotic spindle and sister chromatid mispositioning that persist until telophase, indicate that other molecular mechanisms may coordinate plasma membrane remodelling and mitotic spindle dynamics in polarised epithelial cells. We also reveal an accumulation of cytoplasmic vesicles in non-polarised cells, some of which localise to the cleavage furrow, suggesting that asymmetric membrane elongation may also involve vesicle secretion at the cleavage furrow. Given the important role of organelle dynamics in cell division^34^, further studies will be key to characterise these vesicles and dissect the mechanisms that regulate their trafficking and incorporation to the plasma membrane during mitosis. Remodelling of the plasma membrane in HeLa cells is accompanied with polar cortical blebbing which re-centres the mitotic spindle and releases cortical tensions to stabilise cell shape and ensure correct positioning of the cleavage furrow ^38, 39^. Importantly, our results in non-polarised mammary epithelial cells showing that blebbing does not rescue defects in mitotic spindle, sister chromatids and cleavage furrow positioning, reinforce a model whereby E-cadherin-mediated cell-cell adhesion is crucial for mitotic cells to sense and integrate external forces from neighbouring cells, which in turn decreases intercellular cortical tensions to ensure symmetric cell division. Thus, in contrast to isolated cultured cells, mammary epithelial cells must interact with their tissue environment to maintain their epithelial identity and native geometry to execute error-free mitosis. One must be cautious when modelling and generalising some mechanisms of cell division; cellular identity (in this case, the epithelial origin) defines the response of mitotic cells to certain environmental cues, which may not be observed in non-epithelial cells.

We show that non-polarised mitotic mammary epithelial cells roundup faster and are rounder as compared to polarised cells, further pointing to differences with findings in isolated cells^15^ and the important role of cell-cell adhesion in the regulation of the mechanics of mitosis in epithelial cells. Re-organisation of the actomyosin cytoskeleton into a uniform, contractile cortical meshwork, is a key mechanism for the generation of cortical tensions at the cell surface that drive mitotic rounding, which culminates during metaphase^16, 17^. Remarkably, we found that loss of cell-cell adhesion integrity leads to an increase in the thickness of cortical F-actin in metaphase cells, which correlates with efficient rounding. This contrasts with findings in isolated HeLa cells where the thinning of cortical actin is crucial for generating cortical tensions that drive efficient mitotic rounding^15^. We speculate that cortical actin thickening in non-polarised mitotic mammary epithelial cells acts as a compensatory mechanism that allows cells to overcome loss of tensions and traction forces that polarised cells receive *via* E-cadherin, from neighbouring cells and the epithelial tissue layer. Consistent with this hypothesis, we show that polarised cells have a thinner F-actin cortex and adopt an elongated shape that is defined by neighbouring cells throughout mitosis, where the mitotic spindle aligns along the longest axis of the cell. E-cadherin interacts with actin to sense external mechanical forces and mediate their transduction into intercellular signalling^55–58^. It has been shown that loss of E-cadherin impairs F-actin integrity, which contributes to morphological changes that induce epithelial-to-mesenchymal transition (EMT)^59^. E-cadherin regulates the localisation and function of DDR1 (discoidin domain receptor 1) and the Rho GTPase RhoE, which in turn inhibits ROCK either directly or through RhoA-GTP, leading to a decrease in the actomyosin contractility between individual cells, thereby maintaining proper epithelial integrity^60^. Through a similar mechanism, E-cadherin-mediated decrease in actomyosin contractility controls mitotic spindle assembly and centrosome dynamics^61^. Vinculin has been shown to connect E-cadherin to F-actin at adherens junctions to coordinate mitotic rounding with epithelial integrity maintenance^62^. Interestingly, in our study, fast and efficient rounding-up of non-polarised mammary epithelial cells did not prevent defects in mitotic spindle dynamics or mitotic progression and outcome. This is again different from findings in HeLa cells showing that efficient cell rounding ensures correct mitotic spindle assembly and chromosome segregation^15^. This further points to the crosstalk between mitotic mammary epithelial cells and their neighbouring cells, through E-cadherin, to coordinate correct epithelial integrity with cortical actin organisation and mitotic mechanics. Consistent with this idea, rounding mitotic MDCK cells generate pushing forces against tissue confinement to create space necessary for their division and correct mitotic spindle orientation and chromosome segregation^25^. Reciprocally, maintenance of cell-cell adhesion integrity allows neighbouring cells to exert tensions and traction forces that fine-tune mitotic cell rounding for correct mitosis mechanics and progression^14, 25, 45^.

Correct crosstalk between actin and astral microtubule at the cell cortex is crucial for generating balanced cortical forces that ensure proper assembly and alignment of the mitotic spindle, as well as for coordinating of chromosome segregation with cytokinesis to ensure error-free cell divisions^2, 7, 24^. In addition to the effects on cortical F-actin rearrangement in non-polarised cells, our experiments reveal an increase in the length of astral microtubules, indicating their stabilisation. This results in mitotic spindle misorientation and excessive oscillations which persist during anaphase. The increase in mitotic spindle length in non-polarised cells did not rescue spindle and chromatid centring defects, which is again different from findings in HeLa cells where larger mitotic spindles are centrally placed in cells undergoing asymmetric plasma membrane elongation to ensure the separation of equal-sized daughter cells at cytokinesis^38^. Our results showing several defects in mitotic spindle assembly upon loss of cell-cell adhesion integrity including an increase in the proportion of multipolar spindles, could explain the high incidence of chromosome miss-segregation and delayed mitotic progression. We also propose that our observed defects in mitotic spindle dynamics in non-polarised cells are likely due to the thinning of the subcortical actin clouds, which have been shown to associate with retraction fibres and regulate the growth of astral microtubules and their interaction with cortical actin during mitosis^10, 63, 64^. The vesicle-bound NDP52 protein recruits the actin assembly factor N-WASP to regulate the dynamics of the subcortical actin ring, which in turn acts on the growth of astral microtubules to regulate LGN-mediated mitotic spindle orientation, and precise chromosome segregation^10^. Similarly, the myosin 10 motor protein associates with astral microtubules and works in parallel with Dynein to regulate microtubule dynamic interaction with the cortex and orient centrosomes towards the subcortical actin clouds during mitosis^64^. Recent evidence in several mammalian epithelial cells *in vivo* indicates that cell shape and polarity coordinate the interplay between the cortex, mitotic spindle dynamics and chromosome segregation^22^. Nonetheless, the underlying molecular mechanisms remain poorly characterised. In the mouse prostate epithelium, conditional knockout of E-cadherin affects the LGN-NuMA-Scrib complex resulting in mitotic spindle orientation defects that lead to loss of epithelial integrity that causes carcinogenesis^23^. In MCF-10A cells, E-cadherin-mediated reduction of cortical tensions promotes mitotic spindle bipolarity and acts through myosin 10 to regulate the dynamics and clustering of centrosomes^61^. During planar cell division, astral microtubule plus-ends are coupled to E-cadherin junctions that are associated with the actin cortex^7^. E-cadherin and another junctional protein afadin directly bind LGN to promote planar spindle orientation in MDCK and Caco-2 cells, respectively^4, 12^. E-cadherin senses tensile forces from neighbouring cells at adherens junctions to pattern LGN at the cell cortex^12, 27^, while afadin concomitantly interacts with F-actin and LGN to regulate cortical F-actin reorganisation^4^, thereby promoting polarised cortical recruitment of Dynein and anchoring of astral microtubules. Future studies will be key to characterise actin– and microtubule-associated proteins and crosslinkers that regulate the dynamics of F-actin and astral microtubules and their crosstalk in mitotic mammary epithelial cells and determine how their function is coordinated with cell-cell adhesion.

Here we propose a mechanism whereby the membrane-associated S100A11 protein acts as a molecular sensor of external cues linking plasma membrane remodelling to E-cadherin-dependent cell adhesion to coordinate cortex rearrangement and LGN-mediated mitotic spindle dynamics for correct progression and outcome of mammary epithelial cell divisions. S100A11 is a member of the S100 family of proteins containing two EF-hand Ca^2+^-binding motifs, which promote S100A11 homodimerization and define its functions^65^. S100A11 homodimer interacts with two molecules of ANXA1 to form a heterotetrameric complex that regulates several cellular processes, including connecting phospholipid membranes to mediate vesicular trafficking and recycling, regulation of actin dynamics and its interaction with the plasma membrane^46, 66, 67^. S100A11 acts with ANXA1 and ANXA2 to coordinate plasma membrane remodelling and repair with cortical actin polymerisation, which promotes survival and migration of metastatic cells^47^. In the presence of Ca^2+^, S100A11 binds directly to actin to inhibit actin-activated myosin ATPase activity of smooth muscle cells, indicating a key role of S100A11 in the regulation of actin-myosin-mediated contractility^66^. In mitotic mammary epithelial cells, we recently showed that localisation of S100A11-ANXA1 to the plasma membrane is required for polarised accumulation of LGN at the lateral cortex to promote planar cell division^9^. S100A11 has been shown to associate to microtubules of keratinocytes, which facilitates its translocation to the plasma membrane in a Ca^2+^-dependent manner^67^. Here, we provide a first characterisation of S100A11 dynamic localisation during the cell cycle and reveal that the protein distributes in both the cytoplasm and plasma membrane throughout mitosis. Importantly, we show that S100A11 knockdown phenocopies the effect of cell-cell adhesion perturbation, impairing plasma membrane remodelling and resulting in similar defects in mitotic progression and outcome. Our observations that S100A11 and E-cadherin co-localisation at the cell surface in polarised cells and their co-translocation to the cytoplasm upon loss of cell-cell adhesion further points to a key role of S100A11 in coordinating the interplay between plasma membrane remodelling and E-cadherin function. Consistent with this, we reveal that S100A11 knockdown abrogates the polarised clustering of E-cadherin to the lateral cortex and impairs adherens junction formation, affecting cell shape of mitotic and interphase mammary epithelial cells. Our observations that S100A11 depletion impairs cortical actin and astral microtubule organisation – which results in off-centred mitotic spindles in metaphase – also suggest that S100A11 function in the regulation of plasma membrane remodelling and cell-cell adhesion integrity is crucial for the previously described role of E-cadherin in directing localised re-organisation of F-actin and astral microtubules and their crosstalk at the sites of adherens junctions^7^ to ensure correct assembly and positioning of the mitotic spindle. Given evidence that S100A11 associates with F-actin and microtubules^66, 67^, we cannot rule out a direct effect of S100A11 on cortical actin and astral microtubules. Nonetheless, our immunoprecipitation experiments reveal that S100A11 forms a complex with E-cadherin and LGN in mitotic mammary epithelial cells, which corroborates recent proteomics studies showing that S100A11 co-purifies with LGN and E-cadherin in mammary and MDCK cells, respectively^9, 49^. These findings along with those showing that F-actin directly binds LGN^4^, allow us to speculate that S100A11 may be part of the E-cadherin-F-actin-LGN complex to coordinate the crosstalk between adherens junctions, the cell cortex and mitotic machinery. Interestingly, our observations that loss of polarised localisation of LGN to the lateral cortex in S100A11-depleted cells suggests that S100A11 is required for LGN cortical patterning rather than its cytoplasm-to-cortex recruitment, consistent with the function of ANXA1 in the regulation of mitotic spindle planar orientation^9^. It will be important to further dissect the mitotic functions of the S100A11-ANXA1 complex and its interaction with E-cadherin-F-actin-LGN. Related to this, we show that E-cadherin knockdown abrogates LGN recruitment to the cell cortex and impairs mitotic spindle orientation as described previously^12, 23^, but does not affect the localisation of S100A11 to the plasma membrane, further reinforcing the idea that S100A11 acts upstream of E-cadherin to exert its function in mitosis. Finally, our experiments do not reveal a localisation of S100A11 on the mitotic spindle or chromosomes, indicating that the effect S100A11 depletion on chromosome segregation fidelity is a result of impaired cortical and plasma membrane remodelling, consistent with recent studies linking cortical organisation and cell geometry to faithful chromosome segregation^8, 10, 22^. It is likely that S100A11 participates in the transduction of external cues into intracellular signalling manifested by level of faithfulness in chromosome segregation. Together, our findings identify S100A11 as a membrane-associated molecular landmark that bridges plasma membrane remodelling, E-cadherin clustering to adherens junctions, cortical actin-astral microtubule dynamic crosstalk and LGN mitotic spindle machinery, for correct orientation, progression, and outcome of mammary epithelial cell divisions. Further mechanistic studies will be key to understand how the new function of S100A11 in polarised cell division is spatiotemporally regulated.

In conclusion, our study in mammary epithelial cells reveals an important S100A11-dependent functional, dynamic crosstalk between the plasma membrane, E-cadherin-mediated cell adhesion, cortical cytoskeleton, and LGN-mediated mitotic machinery, to ensure error-free cell divisions and correct shape and positioning of daughter cells. While the plasma membrane is established as a physical interface linking external cues to intracellular signalling during cell division^17^, our findings point to a key role of the plasma membrane as a molecular platform including components such as S100A11 that act as mechanical sensors controlling the mechanochemical crosstalk between extracellular and intracellular signalling for correct execution of polarised cell divisions. Future work will be important to characterise the proteome and lipidome of the plasma membrane to identify factors that bridge plasma membrane remodelling and epithelial polarity, and to dissect how they coordinate these with the mitotic machinery. Remarkably, our experiments in non-transformed mammary epithelial cells devoid of oncogenic triggers, demonstrate that perturbation of cell-cell adhesion integrity is sufficient to induce mitotic and cytoarchitectural defects that are known as major contributors to developmental disorders and carcinogenesis^2, 16, 20^. Consistent with this, loss of E-cadherin in the normal prostate epithelium results in mitotic spindle and cell behaviour defects that lead to carcinogenesis^23^. Chromosome instability and chromosome segregation defects have been linked to EMT^20, 68, 69^. Our findings in non-transformed mammary epithelial cells, suggest that loss of cell-cell adhesion integrity and epithelial identity represent a key initiating event leading to aberrant mitotic spindle dynamics and chromosome segregation that drive epithelial malignant transformation. Further studies using 3D cultures and mouse models will be crucial for elucidating how polarised cell divisions are coordinated with external cues to drive mammary epithelial morphogenesis and carcinogenesis, and for understanding how this is defined by polarised epithelial cell identity.

## Materials and Methods

### Cell lines

MCF-10A cells are spontaneously immortalised, non-transformed human mammary epithelial cells, obtained from ATCC® (#CRL-10317). Cells were cultured in Dulbecco’s Modified Eagle Medium/F12 (DMEM/F12, Invitrogen), supplemented with 5% donor horse serum (Gibco), 20 ng/ml human EGF (Sigma #E9644), 0.5 μg/ml Hydrocortisone (Sigma #H0888), 10 μg/ml insulin (Sigma #I1882), 100 ng/ml cholera toxin (Sigma #C8052), 50 U/ml penicillin and 50 μg/ml streptomycin (Life Technologies) and 500 ng/ml Amphotericin B (Gibco #11510496), at 37 °C in a humidified 5% CO_2_ atmosphere.

HEK293 cells (ATCC® #CRL-3216) provided by Melissa Andrews (University of Southampton), were cultured in DMEM with high glucose, sodium pyruvate and L-glutamine (Gibco), supplemented with 10% foetal bovine serum (FBS, Gibco) and 50 U/ml penicillin and 50 μg/ml streptomycin (Life Technologies), at 37 °C in a humidified 5% CO_2_ atmosphere.

### siRNAs and transfections

Transient knockdown of S100A11 and E-cadherin in MCF-10A cells was performed by transfection of MISSION® Predesigned (Sigma) or custom-made siRNAs, respectively. To knockdown S100A11, the following siRNAs were used: SASI_Hs01_00164495 (si-S100A11 #1) and SASI_Hs01_00164498 (si-S100A11 #2). To knockdown E-cadherin, the following siRNAs were used: si-Ecadherin#1 (sense: 5’-CAUCUUGACUAGGUAUUGUCU-3’; antisense: 5’– AGACAAUACCUAGUCAAGAUG-3’) and si-Ecadherin#2 (sense: 5’– GAGAGAGUUUCCCUACGUAUA-3’; anti-sense: 5’-UAUACGUAGGGAAACUCUCUC-3’). All experiments using siRNAs were carried in the presence of SiGENOME RISC-Free (Dharmacon), used as a negative control for siRNAs experiments (si-Control). MCF-10A cells were transfected with a final concentration of 50 nm of siRNAs diluted in Opti-MEM® Reduced Serum Medium (Gibco) using Lipofectamine^TM^ RNAiMAX (Invitrogen), following the manufacturer’s protocol. Transfected cells were incubated for 48 h or 72h, before they were live-imaged, lysed or fixed and immunoprocessed.

### Retroviral constructs and generation of stable cell lines

Retroviral constructs were used to transduce MCF-10A cells. Lifeact-mCherry was obtained from Addgene (pTK93_Lifeact-mCherry, #46357). pTK14-GFP-S100A11 was cloned as follows: First, the human S100A11 sequence was obtained from the NCBI database (reference sequence: NM-005620.2). Then, the S100A11 cDNA flanked by XhoI and SacII restriction digestion sites sequences were synthesized in a pEX-A128 plasmid (Eurofins Scientific). Second, the synthesized S100A11 was removed from the pEX-A128 plasmid by restriction digestion with XhoI and SacII (New England Biolabs), then cloned into a pTK14-GFP plasmid^9^. Correct insertion was verified by Sanger sequencing.

Generation of stable MCF-10A cell lines was performed using retroviral transduction. Retroviruses were prepared in HEK293 cells by calcium phosphate co-transfection of 10 μg retroviral plasmid, 5 μg packaging plasmid pUMVC (Addgene, Plasmid #8449), 6.5 μg envelope plasmid pCMV-VSV-G (Addgene, Plasmid #8454). Virus particles were collected 48 h after transfection then filtered through a 0.45 μm syringe filter and used to infect MCF-10A cells in the presence of 8 μg/ml polybrene (Sigma). Clones of interest were selected using 1 μg/ml puromycin (Sigma).

### Cell cycle synchronization

MCF-10A cells were treated with 9 μM RO-3306 (Sigma #SML0569), dissolved in DMSO, for 18 h to allow CDK1 inhibition and synchronise cells in G_2_/M phase. To further arrest cells in metaphase, MCF-10A cells were released from the G_2_/M block by washing three times in pre-warmed drug-free medium and incubated in fresh medium for 35 min.

### Cell extracts and immunoblotting

MCF-10A cells were lysed in NP-40 lysis buffer [50 mM Trizma® hydrochloride (Tris-HCl), pH 7.4; 250 mM Sodium Chloride (NaCl); 5 mM Ethylenediaminetetraacetic acid (EDTA); 50 mM Sodium fluoride (NaF); 1 mM Sodium orthovanadate (Na_3_VO_4_); 1% Nonidet P40 (NP40)], supplemented with protease inhibitor cocktail (Sigma, #P2714). Protein concentration of lysates was determined using Pierce™ BCA Protein Assay (Thermo Scientific). Proteins were subjected to sodium dodecyl sulphate polyacrylamide gel electrophoresis (SDS-PAGE) and subsequent Western blot analysis. The following primary antibodies were used: anti-α-tubulin DM1A (0.2 μg / ml, Sigma #T6199), anti-E-cadherin (1:500, Abcam #ab76055), anti-S100A11 (1:500, Proteintech #10237-1-AP), anti-LGN (1:500, Sigma #ABT174), anti-phospho-Histone3 (0.2 μg / ml, Sigma #06-570) and anti-GFP (2 μg/ml, Invitrogen #A-11122). Secondary antibodies conjugated to horseradish peroxidase (Invitrogen, #32430, #32460) were used at 1:10,000. Proteins were visualised using SuperSignal™ West Pico PLUS Chemiluminescent Substrate (ECL) (Thermo Scientific), followed by imaging using a Syngene PXi detection system Scanner (Syngene).

### Immunoprecipitation

Clonal MCF-10A stably expressing pTK14-GFP-S100A11 or pTK14-GFP were plated in 15 cm dishes and washed twice with ice-cold PBS before protein extraction. About 10 × 10^7^ cells were lysed in a mild lysis buffer [50 mM Tris, pH 7.4, 150 mM NaCl, 0.5 mM EDTA, 10 mM NaF, 1 mM Na3VO4, 0.5% Nonidet P40 (NP40)], containing protease inhibitor cocktail. Cell lysates were cleared by centrifugation at 17,000 x *g* for 30 min at 4°C. Co-immunoprecipitation was performed using a GFP-Trap Kit (Chromotek, #gtma-20) following manufacturer’s instructions. For immunoblotting, washed beads were eluted by boiling in Laemmli sample buffer (Bio-Rad) containing 5% 2-mercaptoethanol.

### Immunofluorescence

The following primary antibodies were used: anti-E-cadherin (1:200, Fisher, #13-1900), anti-S100A11 (10 μg/ml, Proteintech #10237-1-AP), anti-LGN (1:200, Sigma #ABT174), anti-α-tubulin DM1A (1 μg/ml, Sigma #T6199), anti-γ-tubulin AK-15 (1:300, Sigma #T3320). AlexaFluor555 phalloidin (1:50, Life Technologies #A34055) was used to label F-actin. Secondary antibodies (Life Technologies) used were goat anti-mouse (#A-32723 and #A-21125), anti-rabbit (#A-11037 and #A-11008) and anti-rat (#A-11007) conjugated to AlexaFluor488, AlexaFluor594 or AlexaFluor647, at 5 μg/ml.

To visualise α-tubulin and γ-tubulin, MCF-10A cells were fixed in ice-cold anhydrous methanol (Sigma) for 5 minutes followed by fixation with 4% paraformaldehyde (PFA, Fisher) for 10 min. PFA was prepared in PHEM [60 mM Piperazine-1,4-bis(2-ethanesulfonic acid) (PIPES), 25 mM 4-(2-hydroxyethyl)-1-piperazineethanesulfonic acid (HEPES), 10 mM ethylene glycol-bis(β-aminoethyl ether)-N,N,N′,N′-tetraacetic acid (EGTA), and 4 mM Magnesium Sulphate (MgSO4)], diluted in PBS. Cells were washed with PBS, then permeabilised with 0.1% Triton X-100-PBS (Sigma) for 10 min. Cells were blocked with 10% goat serum (Sigma) and 1% Bovine Serum Albumin (BSA, Sigma) in 0.1% Triton X-100-PBS for 1 h. To visualise E-cadherin, S100A11 and LGN, cells were fixed in ice-cold anhydrous methanol for 10 minutes. Cells were then washed with PBS and incubated in 10% goat serum and 1% BSA in 0.1% Triton X-100-PBS for 1 h. To visualise cortical F-actin and astral microtubules, cells were fixed 10 min in 3% PFA and 0.25% Glutaraldehyde (Sigma) in 0.2% NP-40 (Sigma) diluted in Brinkley Buffer 1980 (BRB80) [80 mM PIPES, 1 mM Magnesium Chloride (MgCl2), hexahydrate and 1 mM EGTA diluted in dH2O]. Cells were blocked in 0.1% ammonium chloride (NH_4_Cl) (Sigma) in BRB80 for 10 min, followed by two washes in BRB80 for 5 min each. Cells were then blocked in 3% BSA in 0.2% NP-40-BRB80. For all immunostaining, cells were incubated with primary antibodies at 4°C overnight. Cells were washed and incubated with appropriate secondary antibodies for 1 h at RT. Finally, cells were counterstained with 25 µg/ml Hoechst 33342 (Sigma) and mounted with Vectashield antifade mounting medium (Vector Laboratories).

### Quantitative confocal microscopy

Immunofluorescence images were captured with an inverted Leica TCS SP8 inverted laser scanning microscope (Leica Microsystems) using a 63x glycerol immersion (63x HC Plan/Apo CS2 1.30 NA) objective. *Z*-stacks at 16-bit depth and 2048 × 2048 pixels were collected at 0.2 or 0.3 μm intervals. Images were processed with Fiji software^70^. Astral microtubule images were denoised using the MATLAB-based ND-Safir software^71^.

Cortical fluorescence intensity of E-cadherin in metaphase cells was measured in Fiji using a custom macro as described in^8^. The macro measured the pixel value at the cell cortex providing 180 measurements with 2-degree intervals around the defined cell cortex. Fluorescence intensity values were reported along the cortex starting (and finishing) from a point facing the metaphase plate. For measurements along the *xy* axis, position 0 and 180 face the chromosome plate (central cortex) and position 90 and 270 face the spindle poles (lateral cortex): 180 positions were scanned (every 2°) along the cortex. Along the *z* axis, positions 0° and 360° indicate the apical cortex, 90° and 270° indicate the lateral cortex, and 180° indicates the basal cortex. To account for more elongated cell shapes, a line was drawn along the cell contour using the Freehand tool and the macro fits an ellipse to this contour. Fluorescence intensities were calculated along a 30 pixel-long radial line overlapping the cortex and the maximum intensity was reported. Fluorescence was measured around a 10-pixel perpendicular line along the cortex. Measurements were taken at 280 positions, starting from the short axis of the ellipse. Background values were subtracted from all measured fluorescence intensities.

To measure relative fluorescence intensities of E-cadherin and S100A11 at the cell cortex and cytoplasm, a 30-pixel line was drawn across the lateral surface and the cytoplasm using Fiji software. The line scan function of Fiji was used to reveal the relative fluorescence intensity across the line.

Measurement of F-actin fluorescence intensities was performed on 8-bit images, generating a selection for measuring total F-actin fluorescence intensity (*I*_total actin_). Cortical F-actin (*I*_cortical actin_) was measured similarly, with the use of the freehand tool to draw around the cortex of the cell. Subcortical F-actin fluorescence intensity (*I*_subcortical actin_) was determined by subtracting cortical F-actin fluorescence from total F-actin fluorescence. Background values were subtracted for correct F-actin fluorescence intensity measurement on maximum projection images. Actin cloud penetration length was measured by drawing a line toward between the sub-cortical and spindle poles areas at both sides of the cell poles, and then an average value was calculated. Quantification of F-actin organisation at adherens junctions was performed by analysing the clustering of F-actin at sites of cell-cell contacts, which was defined by the formation of F-actin bundles.

Like F-actin, total, plasma membrane and cytoplasmic S100A11 fluorescence intensities were measured. Ratio of plasma membrane-to-cytoplasmic S100A11 fluorescence intensities was measured using Fiji software. Maximum projections of images were generated and the fluorescence intensity of the whole cell (*I*_total_) and the cytoplasm (*I*_cytoplasm_) were measured. Background signal was subtracted, and the plasma membrane-to-cytoplasm fluorescence intensity ratio of S100A11 (*I*_plasma membrane_/*I*_cytoplasm_) was calculated as *I*_plasma membrane_/*I*_cytoplasm_ = (*I*_total_ – *I*_cytoplasm_)/ *I*_cytoplasm_.

Astral microtubule intensity (α-tubulin signal) was measured using Fiji software. Maximum projections of images were generated and the fluorescence intensity of the whole cell (*I*_total_) and the spindle (*I*_spindle_) excluding spindle poles with astral microtubules were measured. Background signal was subtracted, and the relative fluorescence intensity of astral microtubules (*I*_astral, rel_) was calculated as *I*_astral, rel_ = (*I*_total_ – *I*_spindle_)/*I*_spindle_. Length of astral microtubules was measured by drawing a line along the astral microtubule extending towards the cell cortex on both sides of the poles. Similarly, the pole-to-cortex distance was measured by drawing a line towards the closest cell cortex in line with the spindle axis.

Cell segmentation and quantification of cell morphology were performed in cells stained for E-cadherin using the Fiji MorphoLibJ plugin as described in^72^. Cell height was quantified in Fiji by drawing a straight line between the apical and basal membranes of cells viewed along the *z* axis.

Mitotic spindle orientation was measured using Fiji in metaphase cells stained for γ-tubulin. The spindle axis was defined by drawing a 30-pixel wide line across both spindle poles and repositioned along the *z* axis. The spindle axis angle α*z* was measured in respect to the substratum using the angle tool.

### Quantitative live cell imaging

MCF-10A, GFP-S100A11-or Lifeact-mCherry-expressing cells lines were plated in 27 mm Glass Bottom Dishes (Nunc). Prior to imaging, cells were incubated in cell culture medium supplemented with 100 ng/ml Hoechst 33342 (Sigma) for 30 minutes to visualise DNA. When the plasma membrane or mitotic spindles were observed, cells were further incubated in cell culture medium supplemented with CellMask™ Deep Red Plasma Membrane Stain (Thermo Scientific # C10046) at 1:1000 dilution for 10 minutes or with 100 nM SiR-tubulin (Spirochrome #251SC002) for 3 h. Cells were imaged at 37 °C in CO_2_ independent medium (Gibco) using a DeltaVision Elite microscope (GE Healthcare) coupled to a sCMOS max chip area 2048 × 2048 camera (G.E. Healthcare). For each recording, image stacks at 0.6 μm increments in 1,024 x 1,024 format were acquired using a PlanApo 60x/1.42 Oil immersion objective (Olympus) with 2 × 2 binning. Images were taken at 3 stage positions every 3 min for 3 or 5 h. Exposure times were 200 msec and 5% laser power for GFP, 50 msec and 2% laser power for mCherry, 80 msec and 2% laser power for labelled DNA and microtubules, and 50 msec and 2% laser power for labelled plasma membrane using the DAPI-FITC-mCh-Cy5 filter set. Images were deconvolved using the DeltaVision software SoftWoRx and further processed using Fiji.

Plasma membrane dynamic remodelling was visualised by following the distribution CellMask^TM^ labelling at the cell surface in successive timeframes, where it displayed circumferential, unilateral, or bilateral accumulation in mitotic cells. Plasma membrane elongation was quantified by merging images of anaphase and telophase using Fiji software. Then, elongation of the plasma membrane at the cell cortex of the newly formed daughter cells was assessed. Equal membrane elongation of the newly generated cells is considered symmetric elongation, whereas unequal elongation is considered asymmetric elongation. Measurement of daughter cells’ area expressed in μm^2^ was performed using the freehand tool in Fiji to highlight the cortex of daughter cells at cytokinesis. Ratio of daughter cell area was calculated by dividing the largest cell area by the smallest cell area. Chromosome-to-cell cortex distance was measured by drawing a line from the middle of the sister chromatid cluster to the polar cell cortex during telophase using the line tool in Fiji. Ratio of chromosome-to-cell cortex distance was calculated by dividing the longest distance by the shortest distance for each cell at telophase.

Oscillations of the mitotic spindle were calculated from time-lapse videos of cells labelled with SiR-tubulin and Hoechst 33342. Metaphase spindle angles in the *z* (α*z*) and *xy* (α*xy*) planes were manually determined for every frame using Fiji software. Measurements of the spindle angle axis in the *z* plane were performed as described above in fixed cells. Spindle angles were reported as positive values unless spindle poles changed direction. In this case, angles were displayed as negative values. Mitotic spindle orientation in the *xy* plane was measured by drawing a line crossing both spindle poles of the first frame of metaphase. This line was embedded to all the following frames to mark the initial position of the mitotic spindle. In the next frame, another line was overlaid across the poles to define the new spindle axis. The angle between the initial spindle position and the current spindle axis was measured using the angle tool. Spindle movement directions were considered by reporting spindle rotations clockwise as positive angles and reverse movements as negative angles. Spindle angle deviations greater than 10° between two frames were counted as oscillations. The oscillation index was determined as the percentage of oscillations in respect to the total number of frames.

Quantification of actin fluorescence from time-lapse videos of Lifeact-mCherry-expressing cells in prometaphase and metaphase was performed as described in^73^. Ratios of prometaphase-to-metaphase total, actin and subcortical were calculated. Cell morphology and roundness (4*area/pi*sqr(major axis)) quantification was performed using the shape descriptors plugin in Fiji. Rounding timing was determined by analysing successive frames of the time-lapse videos. The full width at half maximum (FWHM) of cortex intensity profiles to assess the thickness of cortical actin. A straight line with the same length was drawn across the cortical actin in all data analysed (cortical actin was centred at the middle of the line). Then, a plot profile was generated using Fiji, and the FWHM calculated using Origin(Pro) Version 2021b (https://www.originlab.com/2021).

### Statistical analysis

All statistical analysis were performed with GraphPad Prism 9.1.2 software. Graphs were created using GraphPad Prism 9.1.2 software and RStudio (1.2.5033 version). Multiple groups were tested using analysis of variance (ANOVA) with post hoc Tukey or Dunnett tests, and comparisons between two groups were performed using *t*-tests. Data are shown as mean ± standard error of the mean (s.e.m.) from three or four independent experiments. *P* ≤ 0.05 was considered statistically significant. Asterisks indicate levels of significance (\**P* ≤ 0.05; \*\**P* ≤ 0.01; \*\*\**P* ≤ 0.001; \*\*\*\**P* ≤ 0.0001).

## Supplementary Information

The online version contains four supplementary Figures and twelve Movies available at

## Acknowledgments

We would like to thank Dr. Mark Willett at the Imaging and Microscopy Centre for valuable assistance with fluorescence microscopy and Dr. Melissa Andrews for providing the HEK293 cell line. Cartoons in the main and supplementary Figures were created using BioRender.com. This work was supported by a Wellcome Trust Seed Award in Science (210077/Z/17/Z) and MRC New Investigator Research Grant (MR/R026610/1) awarded to SE. MMH was supported by a Kingdom of Saudi Arabia Ministry of Education PhD studentship.

## Author Contributions

M.M.H. designed and performed experiments, analysed, and interpreted the data. J.C. performed experiments and analysed the data. M.F. performed experiments. M.P. designed experiments and supervised M.M.H. S.E. conceived and designed the project, performed experiments, analysed, and interpreted the data, supervised M.M.H. and wrote the manuscript with the help of M.M.H. All the authors provided intellectual input, edited, and approved the final manuscript.

---

**Figure S1.**
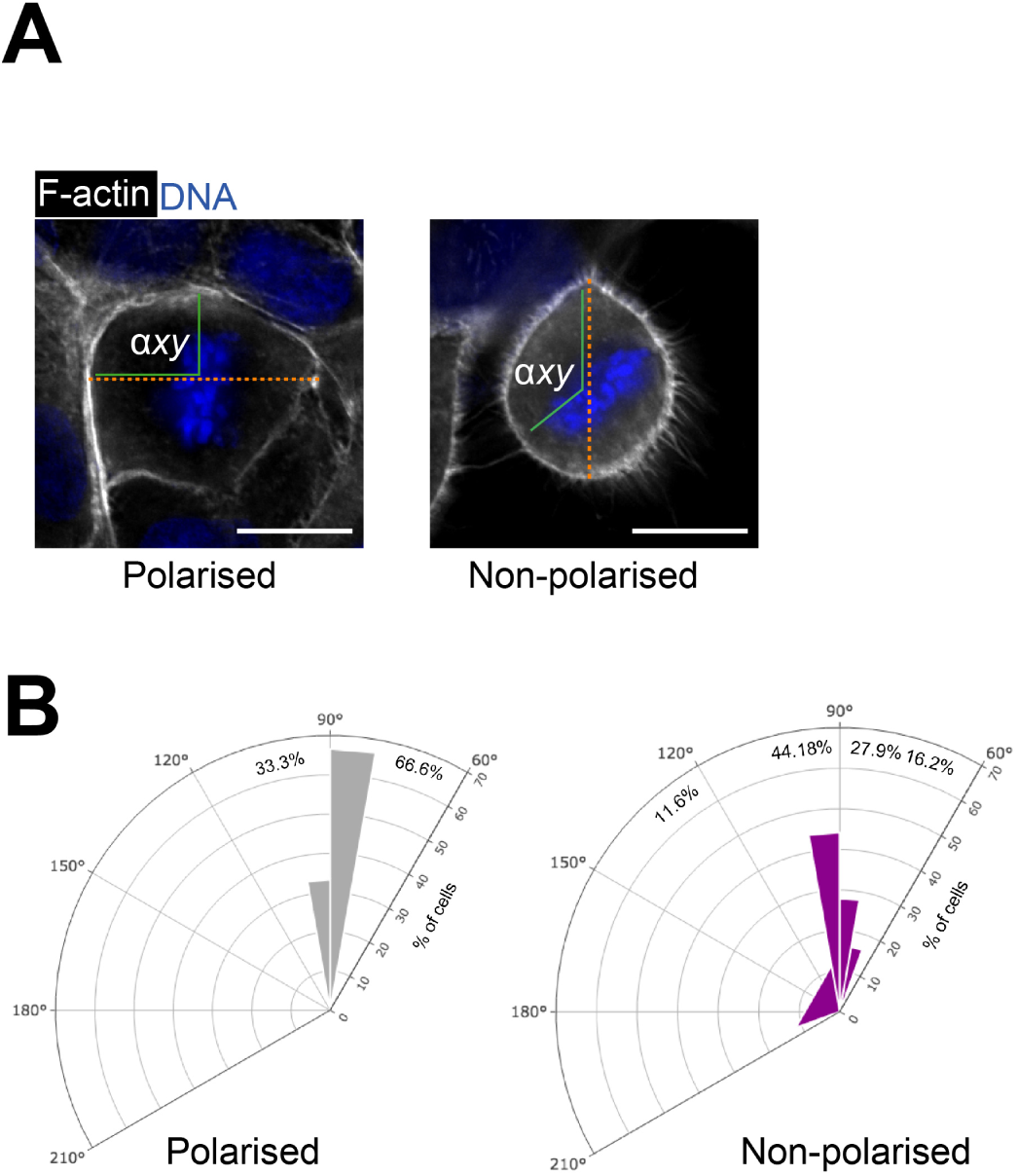
The mitotic spindle aligns following Hertwig’s rule in polarised cells but not in non-polarised cells. (**A**) Confocal images of representative polarised and non-polarised MCF-10A cells stained for F-actin (grey) and counterstained with Hoechst 33342 (DNA, blue). The orange dashed line indicates the long axis of the cell. The angle α*xy* indicates the orientation of the metaphase plate relative to the long axis of the cell. Scale bar, 10 µm. **(B)** α*xy* angle frequencies in polarised and non-polarised cells (polarised: *n* = 45 cells; non-polarised: *n* = 40 cells, 3 independent experiments).

**Figure S2.**
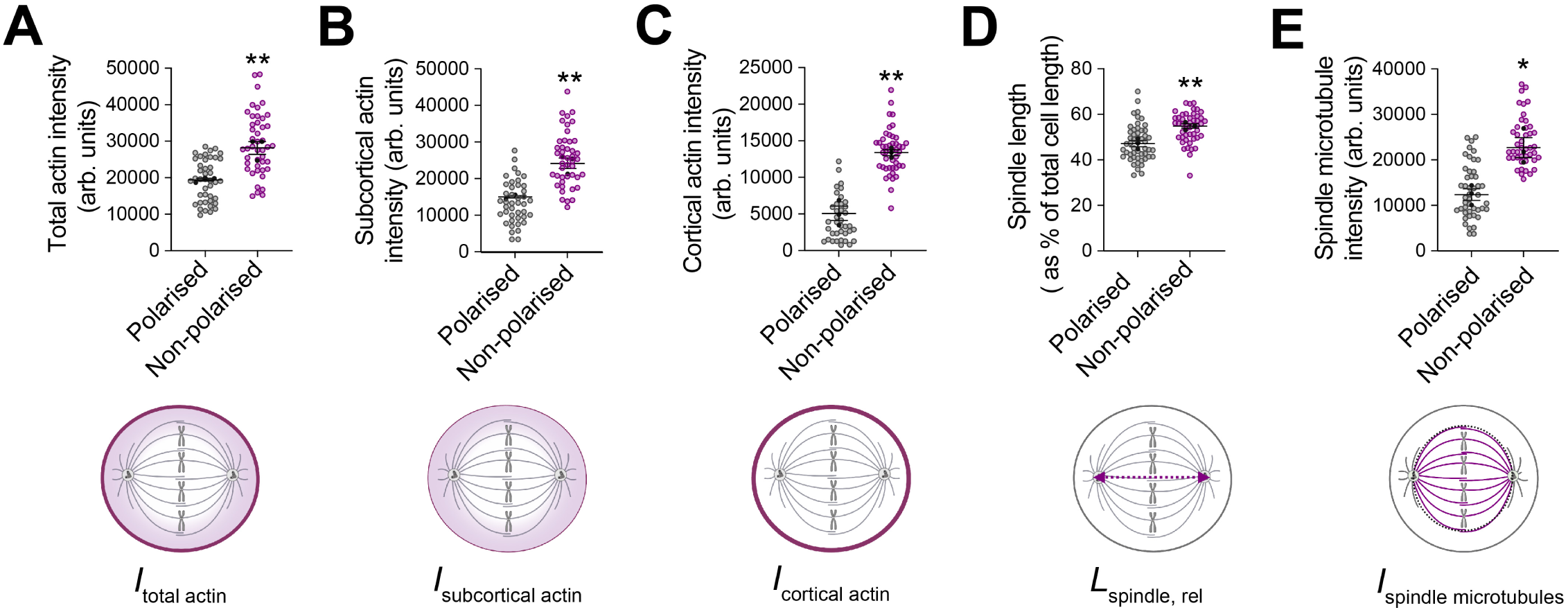
Loss of cell-cell adhesion integrity affects F-actin organisation and mitotic spindle assembly in fixed metaphase cells. (**A-C**) Relative fluorescence intensity of total F-actin, subcortical F-actin, and cortical F-actin in polarised and non-polarised MCF-10A cells (polarised: *n* = 42 cells; non-polarised: *n* = 45 cells). Two-sided *t*-test, total: \*\**P* = 0.0082; subcortical: \*\**P* = 0.0035; cortical: \*\**P* = 0.0015. **(D)** Spindle length in metaphase (polarised: *n* = 51 cells; non-polarised: *n* = 47 cells). Two-sided *t*-test, \*\**P* < 0.0094. **(E)** Relative fluorescence intensity of spindle microtubules (polarised: *n* = 46 cells; non-polarised: *n* = 44 cells). Two-sided *t*-test, \**P* = 0.0147. All data are presented as mean ± s.e.m. from 3 or 4 independent experiments. Source data are provided as a Source Data file.

**Figure S3.**
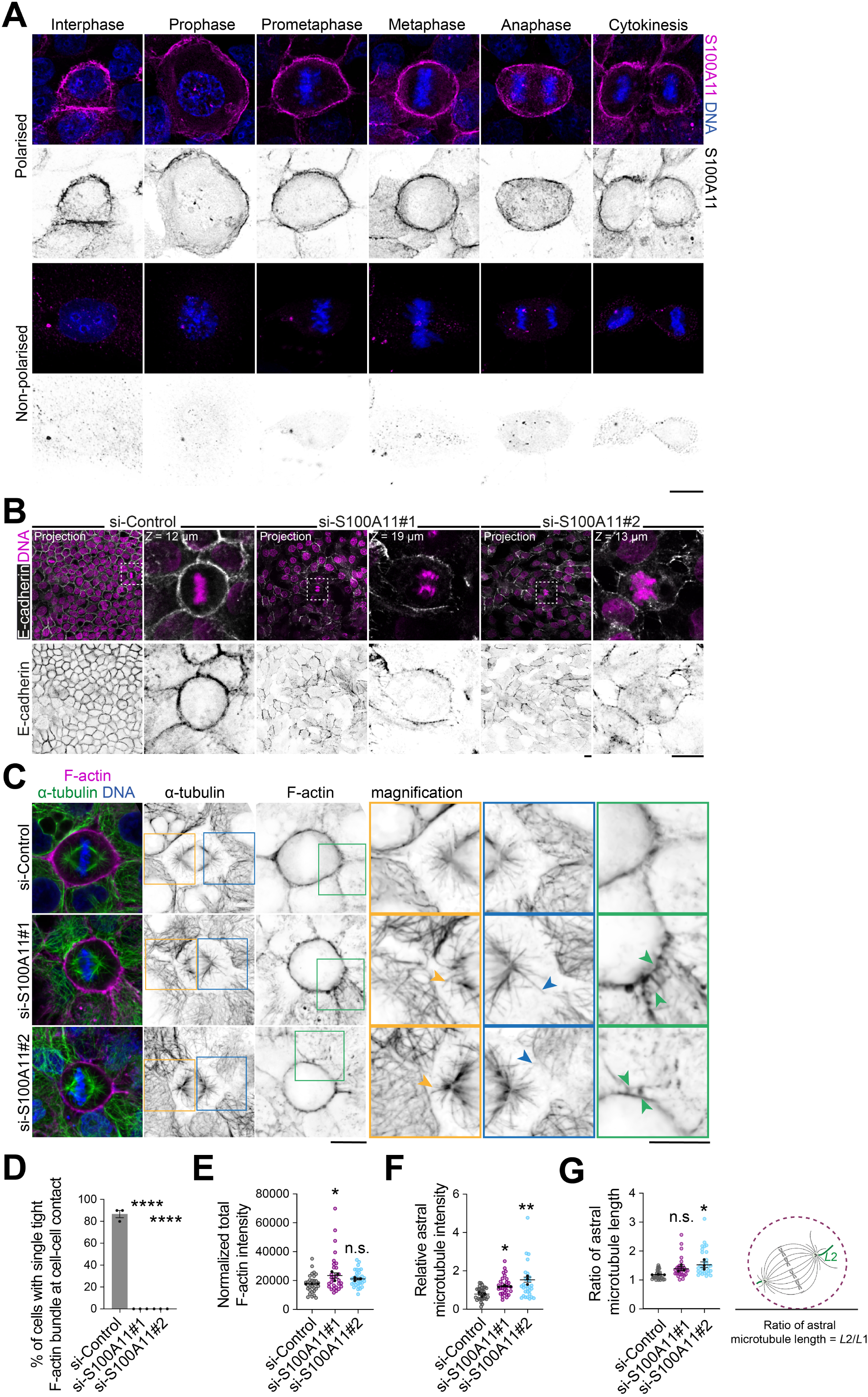
S100A11 distributes at the plasma membrane in polarised cells and regulates F-actin and astral microtubule organisation during mitosis. (**A**) Confocal images of representative polarised and non-polarised MCF-10A cells stained for S100A11 (magenta) and counterstained with Hoechst 33342 (DNA, blue). Scale bar, 10 µm. **(B)** Confocal images of representative si-Control-, si-S100A11#1, si-S100A11#2-treated cells stained for E-cadherin (grey) and counterstained with Hoechst 33342 (DNA, magenta). Scale bar, 10 µm. **(C)** Confocal images of representative si-Control-, si-S100A11#1-, si-S100A11#2-treated metaphase cells stained for α-tubulin (green) and F-actin (magenta), and counterstained with Hoechst 33342 (DNA, blue). Scale bars, 10 µm. **(D)** Percentage of cells with single tight F-actin bundle formation at cell-cell contact in siRNA-transfected cells (si-Control: *n* = 30 cells; si-S100A11#1: *n* = 36 cells; si-S100A11#2: *n* = 38 cells). One-way ANOVA with Dunnett’s test, \*\*\*\**P* < 0.0001; \*\*\*\**P* < 0.0001. **(E)** Relative fluorescence intensity of total actin in siRNA-transfected cells (si-Control: *n* = 32 cells; si-S100A11#1: *n* = 29 cells; si-S100A11#2: *n* = 31 cells). One-way ANOVA with Dunnett’s test, \**P* = 0.019; *P* = 0.185. **(F)** Relative fluorescence intensities of astral microtubules in siRNA-transfected cells (si-Control: *n* = 31 cells; si-S100A11#1: *n* = 30 cells; si-S100A11#2: *n* = 30 cells). One-way ANOVA with Dunnett’s test, \**P* = 0.0251 and \*\**P* = 0.0015. **(G)** Ratio of astral microtubule length in siRNA-transfected cells (si-Control: *n* = 29 cells; si-S100A11#1: *n* = 29 cells; si-S100A11#2: *n* = 29 cells). One-way ANOVA with Dunnett’s test, *P* = 0.130; \**P* = 0.021. All data are presented as mean ± s.e.m. from 3 or 4 independent experiments. n.s. (not significant). Source data are provided as a Source Data file.

**Figure S4.**
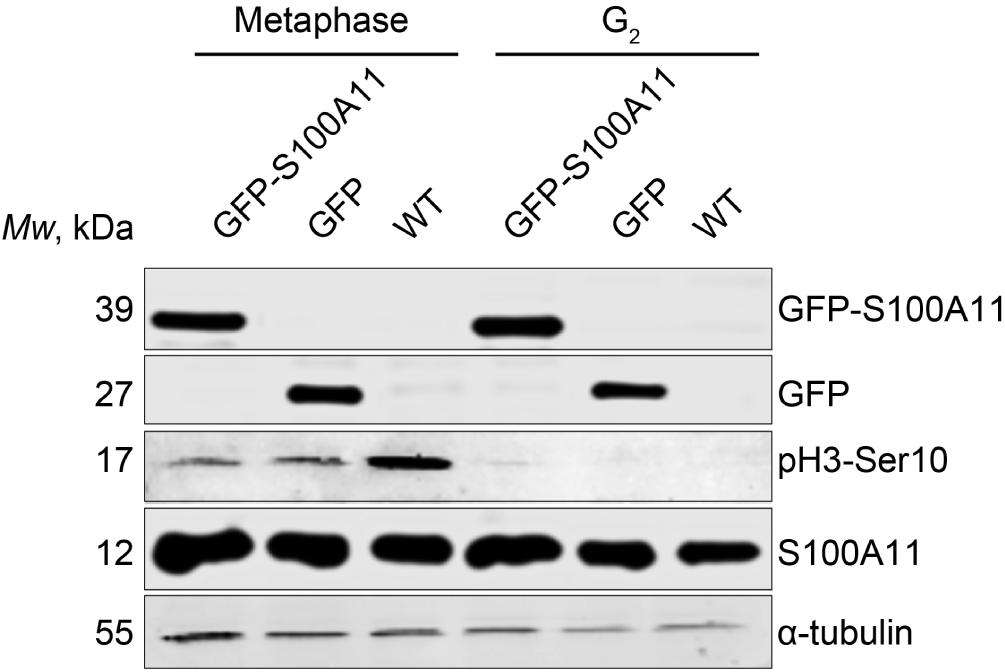
Validation of the cell-cycle synchronisation protocol in mammary epithelial cells. Western blotting of extracts from wild type MCF-10A and clonal MCF-10A stably expressing GFP-S100A11 or GFP cells, synchronised in G_2_ or metaphase. Blots are stained for GFP, S100A11 and phospho-histone H3 (pH3-ser10), and α-tubulin as a loading control (3 independent experiments). Source data are provided as a Source Data file.

## Supplementary Movie description

**Movie S1**

Description: CellMask^TM^ circumferential distribution at the plasma membrane and correct mitosis dynamic progression and outcome in polarised MCF-10A cells. Maximum intensity projections of CellMask^TM^ (green) and Hoechst (DNA, magenta) are shown through time (min).

**Movie S2**

Description: Perturbation of cell-cell adhesion results in unilateral distribution of CellMask^TM^ and asymmetric elongation of the plasma membrane and impairs mitosis dynamic progression and generates unequal-sized daughter cells in non-polarised MCF-10A cells. Maximum intensity projections of CellMask^TM^ (green) and Hoechst (DNA, magenta) are shown through time (min).

**Movie S3**

Description: Perturbation of cell-cell adhesion results in bilateral distribution of CellMask^TM^ and at the plasma membrane and delayed mitosis progression in non-polarised MCF-10A cells. Maximum intensity projections of CellMask^TM^ (green) and Hoechst (DNA, magenta) are shown through time (min).

**Movie S4**

Description: Correct mitotic spindle and chromosome dynamics and mitosis progression and outcome in polarised MCF-10A cells. Maximum intensity projections of SiR-tubulin (green) and Hoechst (DNA, magenta) are shown through time (min).

**Movie S5**

Description: Perturbation of cell-cell adhesion results in chromosome mis-segregation, micronuclei, and delayed mitosis progression in non-polarised MCF-10A cells. Maximum intensity projections of SiR-tubulin (green) and Hoechst (DNA, magenta) are shown through time (min).

**Movie S6**

Description: Perturbation of cell-cell adhesion results in chromosome bridges and delayed mitosis progression in non-polarised MCF-10A cells. Maximum intensity projections of SiR-tubulin (green) and Hoechst (DNA, magenta) are shown through time (min).

**Movie S7**

Description: Perturbation of cell-cell adhesion results in mitotic spindle dynamic and assembly defects, and mitotic arrest in non-polarised MCF-10A cells. Maximum intensity projections of SiR-tubulin (green) and Hoechst (DNA, magenta) are shown through time (min).

**Movie S8**

Description: Correct F-actin dynamic re-organisation and mitotic spindle dynamics and mitosis progression in polarised MCF-10A cells stably expressing Lifeact-mCherry. Maximum intensity projections of Lifeact-mCherry (F-actin, green) and SiR-tubulin (magenta) are shown through time (min).

**Movie S9**

Description: Perturbation of cell-cell adhesion affects F-actin re-organisation and results in mitotic spindle assembly and dynamic defects and impairs mitosis progression and outcome in non-polarised MCF-10A cells stably expressing Lifeact-mCherry. Maximum intensity projections of Lifeact-mCherry (F-actin, green) and SiR-tubulin (magenta) are shown through time (min).

**Movie S10**

Description: S100A11 spatiotemporal distribution in polarised MCF-10A cells stably expressing GFP-S100A11. Maximum intensity projections of GFP-S100A11 (green) and Hoechst (DNA, magenta) are shown through mitosis phases.

**Movie S11**

Description: CellMask^TM^ circumferential distribution at the plasma membrane and correct dynamics of mitosis in polarised MCF-10A cells treated with si-Control. Maximum intensity projections of CellMask^TM^ (green) and Hoechst (DNA, magenta) are shown through time (min).

**Movie S12**

Description: S100A11 knockdown (siS100a11#1) in polarised MCF-10A cells results in unilateral distribution of CellMask^TM^ and asymmetric elongation of the plasma membrane and impairs mitosis dynamic progression and outcome. Maximum intensity projections of CellMask^TM^ (green) and Hoechst (DNA, magenta) are shown through time (min).

## References

1. Prosser, S.L., and Pelletier, L. (2017). Mitotic spindle assembly in animal cells: a fine balancing act. Nat Rev Mol Cell Biol 18, 187–201. 10.1038/nrm.2016.162.

2. Lechler, T., and Mapelli, M. (2021). Spindle positioning and its impact on vertebrate tissue architecture and cell fate. Nat Rev Mol Cell Biol 22, 691–708. 10.1038/s41580-021-00384-4.

3. di Pietro, F., Echard, A., and Morin, X. (2016). Regulation of mitotic spindle orientation: an integrated view. EMBO Rep 17, 1106–1130. 10.15252/embr.201642292.

4. Carminati, M., Gallini, S., Pirovano, L., Alfieri, A., Bisi, S., and Mapelli, M. (2016). Concomitant binding of Afadin to LGN and F-actin directs planar spindle orientation. Nat Struct Mol Biol 23, 155–163. 10.1038/nsmb.3152.

5. Okumura, M., Natsume, T., Kanemaki, M.T., and Kiyomitsu, T. (2018). Dynein-Dynactin-NuMA clusters generate cortical spindle-pulling forces as a multi-arm ensemble. Elife 7. 10.7554/eLife.36559.

6. Pirovano, L., Culurgioni, S., Carminati, M., Alfieri, A., Monzani, S., Cecatiello, V., Gaddoni, C., Rizzelli, F., Foadi, J., Pasqualato, S., and Mapelli, M. (2019). Hexameric NuMA:LGN structures promote multivalent interactions required for planar epithelial divisions. Nat Commun 10, 2208. 10.1038/s41467-019-09999-w.

7. Dogterom, M., and Koenderink, G.H. (2019). Actin-microtubule crosstalk in cell biology. Nat Rev Mol Cell Biol 20, 38–54. 10.1038/s41580-018-0067-1.

8. di Pietro, F., Valon, L., Li, Y., Goiame, R., Genovesio, A., and Morin, X. (2017). An RNAi Screen in a Novel Model of Oriented Divisions Identifies the Actin-Capping Protein Z beta as an Essential Regulator of Spindle Orientation. Curr Biol 27, 2452–2464 e2458. 10.1016/j.cub.2017.06.055.

9. Fankhaenel, M., Hashemi, F.S.G., Mourao, L., Lucas, E., Hosawi, M.M., Skipp, P., Morin, X., Scheele, C., and Elias, S. (2023). Annexin A1 is a polarity cue that directs mitotic spindle orientation during mammalian epithelial morphogenesis. Nat Commun 14, 151. 10.1038/s41467-023-35881-x.

10. Yu, H., Yang, F., Dong, P., Liao, S., Liu, W.R., Zhao, G., Qin, B., Dou, Z., Liu, Z., Liu, W., et al. (2019). NDP52 tunes cortical actin interaction with astral microtubules for accurate spindle orientation. Cell Res 29, 666–679. 10.1038/s41422-019-0189-9.

11. Chiu, C.W.N., Monat, C., Robitaille, M., Lacomme, M., Daulat, A.M., Macleod, G., McNeill, H., Cayouette, M., and Angers, S. (2016). SAPCD2 Controls Spindle Orientation and Asymmetric Divisions by Negatively Regulating the Galphai-LGN-NuMA Ternary Complex. Dev Cell 36, 50–62. 10.1016/j.devcel.2015.12.016.

12. Gloerich, M., Bianchini, J.M., Siemers, K.A., Cohen, D.J., and Nelson, W.J. (2017). Cell division orientation is coupled to cell-cell adhesion by the E-cadherin/LGN complex. Nat Commun 8, 13996. 10.1038/ncomms13996.

13. Matsumura, S., Hamasaki, M., Yamamoto, T., Ebisuya, M., Sato, M., Nishida, E., and Toyoshima, F. (2012). ABL1 regulates spindle orientation in adherent cells and mammalian skin. Nat Commun 3, 626. 10.1038/ncomms1634.

14. Lancaster, O.M., Le Berre, M., Dimitracopoulos, A., Bonazzi, D., Zlotek-Zlotkiewicz, E., Picone, R., Duke, T., Piel, M., and Baum, B. (2013). Mitotic rounding alters cell geometry to ensure efficient bipolar spindle formation. Dev Cell 25, 270–283. 10.1016/j.devcel.2013.03.014.

15. Serres, M.P., Samwer, M., Truong Quang, B.A., Lavoie, G., Perera, U., Gorlich, D., Charras, G., Petronczki, M., Roux, P.P., and Paluch, E.K. (2020). F-Actin Interactome Reveals Vimentin as a Key Regulator of Actin Organization and Cell Mechanics in Mitosis. Dev Cell 52, 210–222 e217. 10.1016/j.devcel.2019.12.011.

16. Taubenberger, A.V., Baum, B., and Matthews, H.K. (2020). The Mechanics of Mitotic Cell Rounding. Front Cell Dev Biol 8, 687. 10.3389/fcell.2020.00687.

17. Rizzelli, F., Malabarba, M.G., Sigismund, S., and Mapelli, M. (2020). The crosstalk between microtubules, actin and membranes shapes cell division. Open Biol 10, 190314. 10.1098/rsob.190314.

18. Leguay, K., Decelle, B., Elkholi, I.E., Bouvier, M., Cote, J.F., and Carreno, S. (2022). Interphase microtubule disassembly is a signaling cue that drives cell rounding at mitotic entry. J Cell Biol 221. 10.1083/jcb.202109065.

19. Matthews, H.K., Delabre, U., Rohn, J.L., Guck, J., Kunda, P., and Baum, B. (2012). Changes in Ect2 localization couple actomyosin-dependent cell shape changes to mitotic progression. Dev Cell 23, 371–383. 10.1016/j.devcel.2012.06.003.

20. Matthews, H.K., Ganguli, S., Plak, K., Taubenberger, A.V., Win, Z., Williamson, M., Piel, M., Guck, J., and Baum, B. (2020). Oncogenic Signaling Alters Cell Shape and Mechanics to Facilitate Cell Division under Confinement. Dev Cell 52, 563–573 e563. 10.1016/j.devcel.2020.01.004.

21. Ramkumar, N., and Baum, B. (2016). Coupling changes in cell shape to chromosome segregation. Nat Rev Mol Cell Biol 17, 511–521. 10.1038/nrm.2016.75.

22. Knouse, K.A., Lopez, K.E., Bachofner, M., and Amon, A. (2018). Chromosome Segregation Fidelity in Epithelia Requires Tissue Architecture. Cell 175, 200–211 e213. 10.1016/j.cell.2018.07.042.

23. Wang, X., Dong, B., Zhang, K., Ji, Z., Cheng, C., Zhao, H., Sheng, Y., Li, X., Fan, L., Xue, W., et al. (2018). E-cadherin bridges cell polarity and spindle orientation to ensure prostate epithelial integrity and prevent carcinogenesis in vivo. PLoS Genet 14, e1007609. 10.1371/journal.pgen.1007609.

24. van Leen, E.V., di Pietro, F., and Bellaiche, Y. (2020). Oriented cell divisions in epithelia: from force generation to force anisotropy by tension, shape and vertices. Curr Opin Cell Biol 62, 9–16. 10.1016/j.ceb.2019.07.013.

25. Sorce, B., Escobedo, C., Toyoda, Y., Stewart, M.P., Cattin, C.J., Newton, R., Banerjee, I., Stettler, A., Roska, B., Eaton, S., et al. (2015). Mitotic cells contract actomyosin cortex and generate pressure to round against or escape epithelial confinement. Nat Commun 6, 8872. 10.1038/ncomms9872.

26. Baker, J., and Garrod, D. (1993). Epithelial cells retain junctions during mitosis. J Cell Sci 104 ( *Pt 2*), 415–425. 10.1242/jcs.104.2.415.

27. Hart, K.C., Tan, J., Siemers, K.A., Sim, J.Y., Pruitt, B.L., Nelson, W.J., and Gloerich, M. (2017). E-cadherin and LGN align epithelial cell divisions with tissue tension independently of cell shape. Proc Natl Acad Sci U S A 114, E5845–E5853. 10.1073/pnas.1701703114.

28. Reinsch, S., and Karsenti, E. (1994). Orientation of spindle axis and distribution of plasma membrane proteins during cell division in polarized MDCKII cells. J Cell Biol 126, 1509–1526. 10.1083/jcb.126.6.1509.

29. Hertwig, O. (1884). Untersuchungen zur morphologie und physiologie der zelle (Fischer).

30. Box, K., Joyce, B.W., and Devenport, D. (2019). Epithelial geometry regulates spindle orientation and progenitor fate during formation of the mammalian epidermis. Elife 8. 10.7554/eLife.47102.

31. Segalen, M., Johnston, C.A., Martin, C.A., Dumortier, J.G., Prehoda, K.E., David, N.B., Doe, C.Q., and Bellaiche, Y. (2010). The Fz-Dsh planar cell polarity pathway induces oriented cell division via Mud/NuMA in Drosophila and zebrafish. Dev Cell 19, 740–752. 10.1016/j.devcel.2010.10.004.

32. Bosveld, F., Markova, O., Guirao, B., Martin, C., Wang, Z., Pierre, A., Balakireva, M., Gaugue, I., Ainslie, A., Christophorou, N., et al. (2016). Epithelial tricellular junctions act as interphase cell shape sensors to orient mitosis. Nature 530, 495–498. 10.1038/nature16970.

33. Nestor-Bergmann, A., Stooke-Vaughan, G.A., Goddard, G.K., Starborg, T., Jensen, O.E., and Woolner, S. (2019). Decoupling the Roles of Cell Shape and Mechanical Stress in Orienting and Cueing Epithelial Mitosis. Cell Rep 26, 2088–2100 e2084. 10.1016/j.celrep.2019.01.102.

34. Carlton, J.G., Jones, H., and Eggert, U.S. (2020). Membrane and organelle dynamics during cell division. Nat Rev Mol Cell Biol 21, 151–166. 10.1038/s41580-019-0208-1.

35. Zhang, X., Bedigian, A.V., Wang, W., and Eggert, U.S. (2012). G protein-coupled receptors participate in cytokinesis. Cytoskeleton (Hoboken) 69, 810–818. 10.1002/cm.21055.

36. Zhang, X., and Eggert, U.S. (2013). Non-traditional roles of G protein-coupled receptors in basic cell biology. Mol Biosyst 9, 586–595. 10.1039/c2mb25429h.

37. Ozlu, N., Qureshi, M.H., Toyoda, Y., Renard, B.Y., Mollaoglu, G., Ozkan, N.E., Bulbul, S., Poser, I., Timm, W., Hyman, A.A., et al. (2015). Quantitative comparison of a human cancer cell surface proteome between interphase and mitosis. EMBO J 34, 251–265. 10.15252/embj.201385162.

38. Kiyomitsu, T., and Cheeseman, I.M. (2013). Cortical dynein and asymmetric membrane elongation coordinately position the spindle in anaphase. Cell 154, 391–402. 10.1016/j.cell.2013.06.010.

39. Sedzinski, J., Biro, M., Oswald, A., Tinevez, J.Y., Salbreux, G., and Paluch, E. (2011). Polar actomyosin contractility destabilizes the position of the cytokinetic furrow. Nature 476, 462–466. 10.1038/nature10286.

40. Uroz, M., Garcia-Puig, A., Tekeli, I., Elosegui-Artola, A., Abenza, J.F., Marin-Llaurado, A., Pujals, S., Conte, V., Albertazzi, L., Roca-Cusachs, P., et al. (2019). Traction forces at the cytokinetic ring regulate cell division and polyploidy in the migrating zebrafish epicardium. Nat Mater 18, 1015–1023. 10.1038/s41563-019-0381-9.

41. Uroz, M., Wistorf, S., Serra-Picamal, X., Conte, V., Sales-Pardo, M., Roca-Cusachs, P., Guimera, R., and Trepat, X. (2018). Regulation of cell cycle progression by cell-cell and cell-matrix forces. Nat Cell Biol 20, 646–654. 10.1038/s41556-018-0107-2.

42. Chaigne, A., Smith, M.B., Lopez Cavestany, R., Hannezo, E., Chalut, K.J., and Paluch, E.K. (2021). Three-dimensional geometry controls division symmetry in stem cell colonies. J Cell Sci 134. 10.1242/jcs.255018.

43. Donker, L., Houtekamer, R., Vliem, M., Sipieter, F., Canever, H., Gomez-Gonzalez, M., Bosch-Padros, M., Pannekoek, W.J., Trepat, X., Borghi, N., and Gloerich, M. (2022). A mechanical G2 checkpoint controls epithelial cell division through E-cadherin-mediated regulation of Wee1-Cdk1. Cell Rep 41, 111475. 10.1016/j.celrep.2022.111475.

44. Lukinavicius, G., Reymond, L., D’Este, E., Masharina, A., Gottfert, F., Ta, H., Guther, A., Fournier, M., Rizzo, S., Waldmann, H., et al. (2014). Fluorogenic probes for live-cell imaging of the cytoskeleton. Nat Methods 11, 731–733. 10.1038/nmeth.2972.

45. Chanet, S., Sharan, R., Khan, Z., and Martin, A.C. (2017). Myosin 2-Induced Mitotic Rounding Enables Columnar Epithelial Cells to Interpret Cortical Spindle Positioning Cues. Curr Biol 27, 3350–3358 e3353. 10.1016/j.cub.2017.09.039.

46. Rintala-Dempsey, A.C., Rezvanpour, A., and Shaw, G.S. (2008). S100-annexin complexes--structural insights. FEBS J 275, 4956–4966. 10.1111/j.1742-4658.2008.06654.x.

47. Jaiswal, J.K., Lauritzen, S.P., Scheffer, L., Sakaguchi, M., Bunkenborg, J., Simon, S.M., Kallunki, T., Jaattela, M., and Nylandsted, J. (2014). S100A11 is required for efficient plasma membrane repair and survival of invasive cancer cells. Nat Commun 5, 3795. 10.1038/ncomms4795.

48. Shankar, J., Messenberg, A., Chan, J., Underhill, T.M., Foster, L.J., and Nabi, I.R. (2010). Pseudopodial actin dynamics control epithelial-mesenchymal transition in metastatic cancer cells. Cancer Res 70, 3780–3790. 10.1158/0008-5472.CAN-09-4439.

49. Guo, Z., Neilson, L.J., Zhong, H., Murray, P.S., Zanivan, S., and Zaidel-Bar, R. (2014). E-cadherin interactome complexity and robustness resolved by quantitative proteomics. Sci Signal 7, rs7. 10.1126/scisignal.2005473.

50. Thery, M., Jimenez-Dalmaroni, A., Racine, V., Bornens, M., and Julicher, F. (2007). Experimental and theoretical study of mitotic spindle orientation. Nature 447, 493–496. 10.1038/nature05786.

51. Dix, C.L., Matthews, H.K., Uroz, M., McLaren, S., Wolf, L., Heatley, N., Win, Z., Almada, P., Henriques, R., Boutros, M., et al. (2018). The Role of Mitotic Cell-Substrate Adhesion Remodeling in Animal Cell Division. Dev Cell 45, 132–145 e133. 10.1016/j.devcel.2018.03.009.

52. Lechler, T., and Fuchs, E. (2005). Asymmetric cell divisions promote stratification and differentiation of mammalian skin. Nature 437, 275–280. 10.1038/nature03922.

53. Taneja, N., Fenix, A.M., Rathbun, L., Millis, B.A., Tyska, M.J., Hehnly, H., and Burnette, D.T. (2016). Focal adhesions control cleavage furrow shape and spindle tilt during mitosis. Sci Rep 6, 29846. 10.1038/srep29846.

54. Ragkousi, K., and Gibson, M.C. (2014). Cell division and the maintenance of epithelial order. J Cell Biol 207, 181–188. 10.1083/jcb.201408044.

55. Buckley, C.D., Tan, J., Anderson, K.L., Hanein, D., Volkmann, N., Weis, W.I., Nelson, W.J., and Dunn, A.R. (2014). Cell adhesion. The minimal cadherin-catenin complex binds to actin filaments under force. Science 346, 1254211. 10.1126/science.1254211.

56. Iskratsch, T., Wolfenson, H., and Sheetz, M.P. (2014). Appreciating force and shape-the rise of mechanotransduction in cell biology. Nat Rev Mol Cell Biol 15, 825–833. 10.1038/nrm3903.

57. Lecuit, T., and Yap, A.S. (2015). E-cadherin junctions as active mechanical integrators in tissue dynamics. Nat Cell Biol 17, 533–539. 10.1038/ncb3136.

58. Yonemura, S., Wada, Y., Watanabe, T., Nagafuchi, A., and Shibata, M. (2010). alpha-Catenin as a tension transducer that induces adherens junction development. Nat Cell Biol 12, 533–542. 10.1038/ncb2055.

59. Chen, A., Beetham, H., Black, M.A., Priya, R., Telford, B.J., Guest, J., Wiggins, G.A., Godwin, T.D., Yap, A.S., and Guilford, P.J. (2014). E-cadherin loss alters cytoskeletal organization and adhesion in non-malignant breast cells but is insufficient to induce an epithelial-mesenchymal transition. BMC Cancer 14, 552. 10.1186/1471-2407-14-552.

60. Hidalgo-Carcedo, C., Hooper, S., Chaudhry, S.I., Williamson, P., Harrington, K., Leitinger, B., and Sahai, E. (2011). Collective cell migration requires suppression of actomyosin at cell-cell contacts mediated by DDR1 and the cell polarity regulators Par3 and Par6. Nat Cell Biol 13, 49–58. 10.1038/ncb2133.

61. Rhys, A.D., Monteiro, P., Smith, C., Vaghela, M., Arnandis, T., Kato, T., Leitinger, B., Sahai, E., McAinsh, A., Charras, G., and Godinho, S.A. (2018). Loss of E-cadherin provides tolerance to centrosome amplification in epithelial cancer cells. J Cell Biol 217, 195–209. 10.1083/jcb.201704102.

62. Monster, J.L., Donker, L., Vliem, M.J., Win, Z., Matthews, H.K., Cheah, J.S., Yamada, S., de Rooij, J., Baum, B., and Gloerich, M. (2021). An asymmetric junctional mechanoresponse coordinates mitotic rounding with epithelial integrity. J Cell Biol 220. 10.1083/jcb.202001042.

63. Inoue, D., Obino, D., Pineau, J., Farina, F., Gaillard, J., Guerin, C., Blanchoin, L., Lennon-Dumenil, A.M., and Thery, M. (2019). Actin filaments regulate microtubule growth at the centrosome. EMBO J 38. 10.15252/embj.201899630.

64. Kwon, M., Bagonis, M., Danuser, G., and Pellman, D. (2015). Direct Microtubule-Binding by Myosin-10 Orients Centrosomes toward Retraction Fibers and Subcortical Actin Clouds. Dev Cell 34, 323–337. 10.1016/j.devcel.2015.06.013.

65. Fritz, G., Botelho, H.M., Morozova-Roche, L.A., and Gomes, C.M. (2010). Natural and amyloid self-assembly of S100 proteins: structural basis of functional diversity. FEBS J 277, 4578–4590. 10.1111/j.1742-4658.2010.07887.x.

66. Zhao, X.Q., Naka, M., Muneyuki, M., and Tanaka, T. (2000). Ca(2+)-dependent inhibition of actin-activated myosin ATPase activity by S100C (S100A11), a novel member of the S100 protein family. Biochem Biophys Res Commun 267, 77–79. 10.1006/bbrc.1999.1918.

67. Eckert, R.L., Broome, A.M., Ruse, M., Robinson, N., Ryan, D., and Lee, K. (2004). S100 proteins in the epidermis. J Invest Dermatol 123, 23–33. 10.1111/j.0022-202X.2004.22719.x.

68. Comaills, V., Kabeche, L., Morris, R., Buisson, R., Yu, M., Madden, M.W., LiCausi, J.A., Boukhali, M., Tajima, K., Pan, S., et al. (2016). Genomic Instability Is Induced by Persistent Proliferation of Cells Undergoing Epithelial-to-Mesenchymal Transition. Cell Rep 17, 2632–2647. 10.1016/j.celrep.2016.11.022.

69. Roschke, A.V., Glebov, O.K., Lababidi, S., Gehlhaus, K.S., Weinstein, J.N., and Kirsch, I.R. (2008). Chromosomal instability is associated with higher expression of genes implicated in epithelial-mesenchymal transition, cancer invasiveness, and metastasis and with lower expression of genes involved in cell cycle checkpoints, DNA repair, and chromatin maintenance. Neoplasia 10, 1222–1230. 10.1593/neo.08682.

70. Schindelin, J., Arganda-Carreras, I., Frise, E., Kaynig, V., Longair, M., Pietzsch, T., Preibisch, S., Rueden, C., Saalfeld, S., Schmid, B., et al. (2012). Fiji: an open-source platform for biological-image analysis. Nat Methods 9, 676–682. 10.1038/nmeth.2019.

71. Boulanger, J., Kervrann, C., Bouthemy, P., Elbau, P., Sibarita, J.B., and Salamero, J. (2010). Patch-based nonlocal functional for denoising fluorescence microscopy image sequences. IEEE Trans Med Imaging 29, 442–454. 10.1109/TMI.2009.2033991.

72. Legland, D., Arganda-Carreras, I., and Andrey, P. (2016). MorphoLibJ: integrated library and plugins for mathematical morphology with ImageJ. Bioinformatics 32, 3532–3534. 10.1093/bioinformatics/btw413.

73. Mahlandt, E.K., and Goedhart, J. (2022). Visualizing and Quantifying Data from Time-Lapse Imaging Experiments. In Fluorescent Microscopy, (Springer), pp. 329–348.

